# Regulating peroxisome-ER contacts via the ACBD5-VAPB tether by FFAT motif phosphorylation and GSK3β

**DOI:** 10.1101/2021.11.11.467785

**Authors:** Suzan Kors, Christian Hacker, Chloe Bolton, Renate Maier, Lena Reimann, Emily J.A. Kitchener, Bettina Warscheid, Joseph L. Costello, Michael Schrader

**Affiliations:** College of Life and Environmental Sciences, Biosciences, University of Exeter, EX4 4QD Exeter, UK; Institute of Biology II, Biochemistry and Functional Proteomics, Faculty of Biology, University of Freiburg, 79104 Freiburg, Germany; CIBSS Centre for Integrative Biological Signalling Studies, University of Freiburg, 79104 Freiburg, Germany

**Keywords:** peroxisomes, membrane contact sites, FFAT motif, phosphorylation, GSK3β

## Abstract

Peroxisomes and the endoplasmic reticulum (ER) cooperate in cellular lipid metabolism. They form membrane contacts through interaction of the peroxisomal membrane protein ACBD5 [acyl-coenzyme A-binding domain protein 5] and the ER-resident protein VAPB [vesicle-associated membrane protein-associated protein B]. ACBD5 binds to the major sperm protein domain of VAPB via its FFAT-like [two phenylalanines (FF) in an acidic tract] motif. However, molecular mechanisms, which regulate formation of these membrane contact sites, are unknown. Here, we reveal that peroxisome-ER associations via the ACBD5-VAPB tether are regulated by phosphorylation. We show that ACBD5-VAPB binding is phosphatase-sensitive and identify phosphorylation sites in the flanking regions and core of the FFAT-like motif, which alter interaction with VAPB and thus, peroxisome-ER contact sites differently. Moreover, we demonstrate that GSK3β [glycogen synthase kinase-3 beta] regulates this interaction. Our findings reveal for the first time a molecular mechanism for the regulation of peroxisome-ER contacts in mammalian cells and expand the current model of FFAT motifs and VAP interaction.

**SUMMARY:** Kors et al. reveal that peroxisome-ER associations via the ACBD5-VAPB tether are regulated by phosphorylation and GSK3β in mammalian cells. Phosphorylation sites in the FFAT-like motif of ACBD5 affect the binding to VAPB and thus, peroxisome-ER contact sites, differently.

## Introduction

Peroxisomes are small, single membrane bound organelles with key roles in cellular lipid and hydrogen peroxide metabolism. They contribute to a wide range of metabolic processes including the β-oxidation of fatty acids and the synthesis of bile acids and plasmalogens (myelin sheath lipids) (Wanders and Waterham, 2006). To fulfil those functions, peroxisomes interact and cooperate with other organelles, such as the endoplasmic reticulum (ER), mitochondria, lysosomes and lipid droplets, in order to efficiently transfer lipid metabolites (e.g. plasmalogen intermediates, chain-shortened acyl-CoAs, cholesterol and very long chain fatty acids (VLCFAs), respectively) (Chu et al., 2015; Wanders et al., 2016; Shai et al., 2018; Chang et al., 2019).

This collaboration requires close proximity of the organelles, which is mediated by protein tethering complexes that physically bridge apposing organelles (Prinz et al., 2020; Silva et al., 2020). Recently, we and others identified novel tethering complexes that mediate membrane contacts between peroxisomes and the ER in mammalian cells (Costello et al., 2017b; c; Hua et al., 2017; Xiao et al., 2019; Guillén-Samander et al., 2021). We revealed that the peroxisomal acyl-CoA binding domain proteins ACBD4 and ACBD5 interact with ER-resident VAMP-associated proteins (VAPs), a protein family widely involved in tethering the ER to other organelles (Murphy and Levine, 2016). This interaction involves a FFAT [two phenylalanines (FF) in an acidic tract] -like motif within ACBD4/5 and the VAP MSP [major sperm protein] domain (Costello et al., 2017b; Hua et al., 2017). Peroxisome-ER contacts control peroxisome movement and positioning, the delivery of lipids required for peroxisomal membrane expansion prior to division and the transfer of lipid metabolites such as cholesterol, plasmalogens and VLCFAs, although the latter has yet to be formally demonstrated (Darwisch et al., 2020; Schrader et al., 2020).

In general, although the overall pattern of membrane contact sites between organelles has been shown to be relatively stable, individual organelle contacts are dynamic (Valm et al., 2017), suggesting that protein tethers between organelles are highly regulated. The importance of dynamism in organelle contacts is exemplified by the mitochondria-ER-cortex tether in yeast which needs to be re-modelled during meiotic divisions (Sawyer et al., 2019). This is achieved by rapid degradation of the organelle tethering complex allowing mitochondrial detachment. Peroxisome-ER contacts also need to be dynamic in order to modulate peroxisome movement and positioning, as well as metabolite flow between the organelles. However, knowledge on how the majority of membrane contact sites are regulated is limited. Previous studies found that several proteins present at membrane contact sites can be regulated by phosphorylation, including lipid transfer proteins CERT, OSBP and ORP3 (Goto et al., 2012; Kumagai et al., 2014; Weber-Boyvat et al., 2015). These proteins have in common that they bind to the MSP domain of VAP via their FFAT motif. This motif consists of the core consensus sequence ^1^EFFDA-E^7^ flanked by a stretch of acidic residues (Loewen et al., 2003). Numerous proteins which possess potential FFAT motifs have now been identified (Slee and Levine, 2019), and previous work has implicated phosphorylation of particular residues in the FFAT motif in regulating VAP binding (Kumagai et al., 2014; Kirmiz et al., 2018; Johnson et al., 2018). Recent studies revealed that VAP can also bind to unconventional FFAT motifs with a serine/threonine residue at position 4, but only when this residue is phosphorylated (phospho-FFAT) to mimic the acidic amino acid present in the conventional motif (Kirmiz et al., 2018; Di Mattia et al., 2020; Guillén-Samander et al., 2021). However, a thorough understanding of the regulation of FFAT-VAP interactions by phosphorylation and how this is linked to kinases and phosphatases as well as other signalling networks is still lacking (Mikitova and Levine, 2012; Murphy and Levine, 2016).

Here, we reveal that peroxisome-ER associations via the ACBD5-VAPB tether are regulated by phosphorylation. We show that the ACBD5-VAPB interaction is phosphatase-sensitive and that ACBD5 phosphomimetic and non-phosphorylatable mutants influence interaction with VAPB and thus, peroxisome-ER contact sites differently. Moreover, we show that glycogen synthase kinase-3 beta (GSK3β) associates with the ACBD5-VAPB complex to regulate peroxisome-ER contacts. GSK3β can directly phosphorylate ACBD5 in the FFAT core at serine-269 (S269), a residue that, when phosphorylated, blocks binding to VAPB. Our findings reveal for the first time a molecular mechanism for the regulation of peroxisome-ER contacts in mammalian cells, provide one of the first clear examples for a physiological role of phosphorylation of peroxisomal proteins in mammals, and expand the current model of FFAT motifs and VAP interaction.

## Results

### The overall phosphorylation status of ACBD5, but not ACBD4, affects VAPB binding

Several studies have suggested a general role for phosphorylation in regulating FFAT-VAP interactions, and multiple phosphorylation sites in ACBD4 and ACBD5 have been reported in high-throughput studies (Bian et al., 2014; Sharma et al., 2014). To confirm ACBD4/5 phosphorylation and to study its role in the interaction between ACBD4 and ACBD5 with VAPB, we first investigated whether ACBD4 and ACBD5 were phosphorylated using the Phos-tag system (Kinoshita et al., 2006). Altered migration in Phos-tag SDS-PAGE following phosphatase treatment indicated that both ACBD4 and ACBD5 were phosphorylated (**Fig. 1A**) as suggested (Bian et al., 2014; Sharma et al., 2014). Next, we tested whether phosphorylation of ACBD4 and ACBD5 had an effect on VAPB binding. COS-7 cells expressing FLAG-ACBD4 or FLAG-ACBD5 were lysed in the presence of phosphatase inhibitor, or lysates were treated with λ protein phosphatase (λPP) (as in **Fig. 1A**). The samples were then incubated with recombinant GST-VAPB, expressed and purified from *E. coli*. Interaction of GST-VAPB with FLAG-ACBD4 was similar in phosphatase inhibitor and phosphatase-treated, immunoprecipitated samples, indicating that the ACBD4-VAPB binding is insensitive to phosphatase treatment (**Fig. 1B**). In contrast, λPP-treated FLAG-ACBD5 failed to bind GST-VAPB (**Fig. 1B**). A FFAT-mutant (mFFAT) of FLAG-ACBD5, which was previously shown to abolish VAPB interaction, was used as a negative control (Costello et al., 2017b). A FFAT mutation in ACBD4 (ACBD4 mFFAT) also resulted in loss of association with GST-VAPB, confirming that, similar to ACBD5, a single FFAT-like motif in ACBD4 is essential for VAPB binding (**Fig. 1B**). Similar results were obtained with endogenous VAPB in a binding assay using lysates of COS-7 cells expressing tagged versions of ACBD4 or ACBD5 (**Fig. 1C, Fig. S1A**). Binding of Myc-ACBD5 to endogenous VAPB was significantly reduced upon λPP-treatment (**Fig. 1C**). We conclude that the interaction between ACBD5 and VAPB, but not ACBD4-VAPB, is highly sensitive to phosphatase treatment.

**Figure 1.**
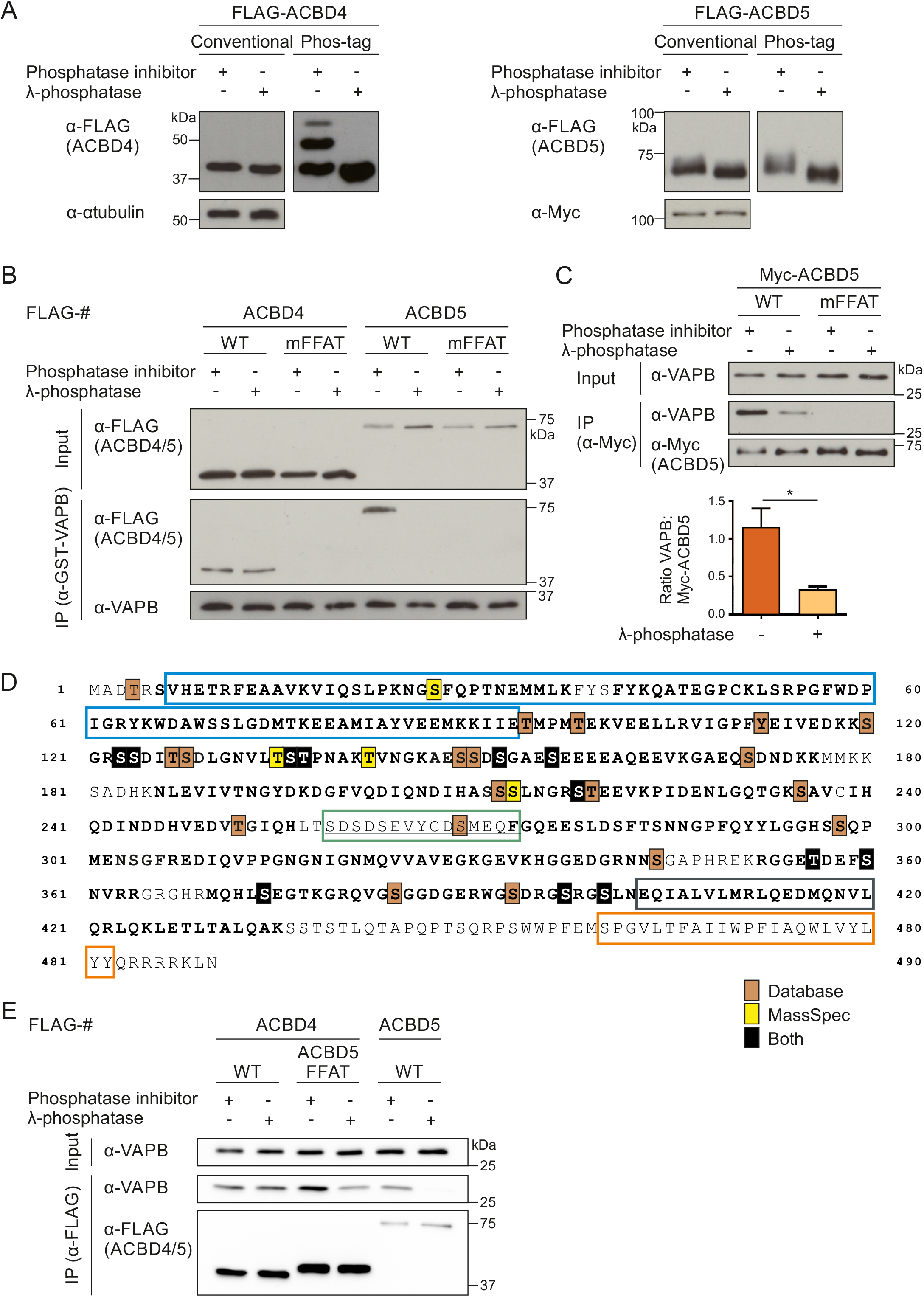
The interaction between ACBD5 and VAPB is sensitive to phosphatase treatment. (**A**) Immunoblots of lysates of FLAG-ACBD4 or FLAG-ACBD5 protein expressed in COS-7 cells treated with or without λ-phosphatase (λPP), using conventional and Phos-tag SDS-PAGE. αTubulin and Myc (unspecific band) served as loading control. (**B**) Binding assay with FLAG-ACBD4 or FLAG-ACBD5 expressed in COS-7 cells and recombinant GST-VAPB. FLAG-ACBD4/5 was treated with λPP. Constructs with mutations in the FFAT-like motif (mFFAT) were used as a negative control. Samples were immunoprecipitated (IP) (GST-TRAP) and immunoblotted using FLAG/VAPB antibodies. (**C**) Myc-ACBD5 was expressed in COS-7 cells, and lysates treated with or without λPP. Myc-ACBD5 was immunoprecipitated and endogenous bound VAPB detected by immunoblotting using Myc/VAPB antibodies. Data were analyzed by a two-tailed unpaired t-test; *, P < 0.05. Error bars represent SEM, with five independent experiments. Total VAPB (IP fraction) was normalized against Myc-ACBD5 (IP fraction). (**D**) ACBD5 protein sequence. Phosphorylation sites identified by database search (Hornbeck et al., 2015; Ullah et al., 2016) and our own mass spectrometry (MS)-based analyses are indicated (DOI: 10.6019/PXD018005) (filled boxes). Coloured boxes enclosing sequence regions indicate different protein domains (colours as in **Fig. 2A**). Bold regions represent peptides identified by MS (-TiO2 and +TiO2). The FFAT-like motif is underlined. (**E**) The FFAT-like motif region of ACBD4 was replaced with that of ACBD5 (ACBD5 FFAT). FLAG-ACBD4/5 constructs were expressed in COS-7 cells and immunoprecipitated to detect endogenous bound VAPB using FLAG/VAPB antibodies. WT, wild type.

**Figure 2.**
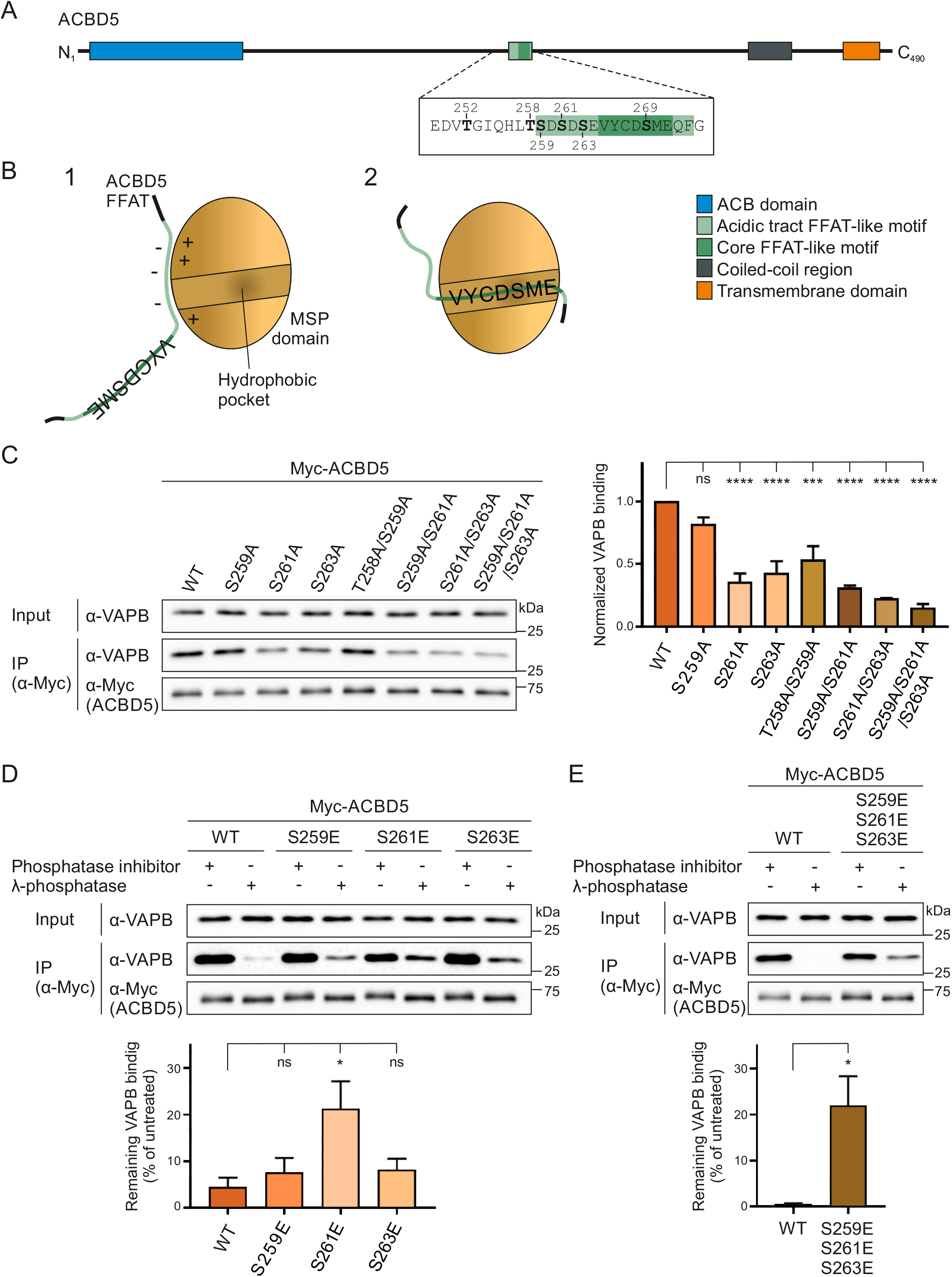
Phospho-mutants of the acidic tract alter the ACBD5-VAPB interaction and its phosphatase-sensitivity. (**A**) Schematic overview of ACBD5 domain structure, including the amino acid sequences of the FFAT-like motif region, with the phosphorylation sites mutated in this study in bold. (**B**) Schematic model of the interaction between the ACBD5 FFAT-like motif and the VAPB MSP domain. The interaction occurs in two steps (Furuita et al., 2010): (1) An initial non-specific electrostatic interaction between the acidic tract of the FFAT motif and the basic electropositive surface of the MSP domain; (2) A specific binding of the FFAT core region to the FFAT-binding site of the MSP domain, which consists of an electropositive face. (**C, D, E**) ACBD5 constructs with non-phosphorylatable (S → A) and phosphomimetic (S → E) residues in the acidic tract were generated and expressed in COS-7 cells. The proteins were immunoprecipitated and endogenous bound VAPB detected by immunoblotting using Myc/VAPB antibodies. (**C**) Data were analyzed by one-way analysis of variance with Dunnett’s multiple comparison test. Total VAPB (IP fraction) was normalized against total VAPB (input) and Myc-ACBD5 (IP fraction). (**D, E**) Lysates were treated with or without λPP before immunoprecipitation. Data were analyzed by one-way analysis of variance with Dunnett’s multiple comparison test (D) or a two-tailed unpaired t-test (E). Total VAPB (IP fraction) was normalized against Myc-ACBD5 (IP fraction). Normalized VAPB signal in the treated sample was then calculated as a percentage of normalized VAPB signal in the untreated sample. ns, not significant; *, P < 0.05; ***, P < 0.001; ****, P < 0.0001. Results of at least three independent IPs were quantified. Error bars represent SEM. WT, wild type.

### ACBD5 phosphorylation profile

To determine the potential phosphorylation sites that are responsible for the phosphatase sensitivity of the ACBD5-VAPB interaction we combined a database search (Hornbeck et al., 2015; Ullah et al., 2016) with our own phosphorylation analysis of ACBD5 by mass spectrometry (MS) (**Fig. 1D**). Multiple phosphorylation sites were identified, but an initial mutational analysis of prominent phosphorylation sites in ACBD5 did not identify any residues involved in VAPB binding (**Fig. S1B**). A more detailed analysis of our MS phosphorylation data revealed that interestingly, there was an apparent gap in the peptide coverage of ACBD5, which included the FFAT-like motif and surrounding region (**Fig. 1D**). As this stretch contained multiple serine/threonine residues and the VAPB binding site, we decided to explore this region by placing it in ACBD4 (**Fig. S2A**). Replacing the FFAT-like motif of ACBD4 with that of ACBD5 now rendered the interaction between ACBD4 and VAPB sensitive to λPP-treatment (**Fig. 1E**). This suggests that phosphorylation in and around the FFAT-like motif of ACBD5 is responsible for the sensitivity of the ACBD5-VAPB interaction to phosphatase treatment. To investigate how phosphorylation of this region can affect VAPB binding, we next generated phosphosite mutants of the highly conserved residues (**Fig. 2A, Fig. S2B-D**).

### Non-phosphorylatable residues in the acidic tract reduce ACBD5-VAPB binding

The core FFAT motif (^1^EFFDA-E^7^) is often surrounded by acidic residues (acidic tract herein), which are thought to contribute to the initial interaction with VAPB as part of a two-step binding model (**Fig. 2B**). The presence of serine/threonine residues in the acidic tract of FFAT(-like) motifs is a common feature (Loewen et al., 2003; Murphy and Levine, 2016), including that of ACBD5 (**Fig. 2A**). Phosphorylation of these residues could mimic canonical aspartic acid and glutamic acid residues, aiding the acidic environment and potentially increasing binding to VAPB. To test this, we generated Myc-tagged single, double and triple ACBD5 phosphosite mutants of the acidic tract by mutating threonine-258 (T258), serine-259 (S259), S261 and S263 to alanine (A), to mimic non-phosphorylated forms. All phospho-mutants were properly targeted to peroxisomes (**Fig. S3A**), as the peroxisome targeting signal located in the transmembrane domain and tail of ACBD4/5 was not altered (Costello et al., 2017a).

The ACBD5 phosphosite mutants were expressed in COS-7 cells and interaction with endogenous VAPB was assessed by immunoprecipitation (**Fig. 2C**). Whereas the S259A single mutant did not significantly reduce VAPB binding, the S261A and S263A single mutants showed a significant reduction, suggesting that phosphorylation of these serine residues in the acidic tract is important for VAPB interaction (**Fig. 2C**). The double mutants T258A/S259A and S259A/S261A also caused a reduction in VAPB binding when compared to WT control, whereas the S261A/S263A double and S259A/S261A/S263A triple mutants had the most prominent effect. This suggests that (i) residues S261 and S263 are the most significant for VAPB binding, and (ii) the overall acidity of the acidic tract contributes to VAPB interaction.

### The acidic tract of ACBD5 contributes to the phosphatase-sensitivity of the ACBD5-VAPB interaction

Overall, our results indicate that lack of phosphorylation of residues in the acidic tract of ACBD5 reduces its binding to VAPB. Therefore, we hypothesized that phosphomimetic mutation of these residues would overcome the sensitivity of the ACBD5-VAPB interaction for phosphatase treatment (**Fig. 1C**). We generated Myc-ACBD5 single and triple site phosphomimetic mutants by replacing S259, S261 and S263 with glutamic acid (E), and assessed their binding to endogenous VAPB by immunoprecipitation after λPP-treatment of the cell lysates (**Fig. 2D, E**). We observed that binding of the single mutants S259E and S263E to VAPB was still reduced upon λPP-treatment, comparable to WT control (**Fig. 2D**). However, the single S261E and triple S259E/S261E/S263E phosphomimetic mutants showed significantly more binding to VAPB than wild type following λPP-treatment (**Fig. 2D, E**). This confirms that phosphorylation of the acidic tract contributes to VAPB binding, but also indicates that additional, as yet uncharacterised phosphatase-sensitive elements exist which mediate VAPB binding.

### Additional phospho-mutants of the FFAT-like motif region affect VAPB binding

ACBD5 contains additional serine/threonine residues further upstream of the FFAT-like motif and within the FFAT core, which have been found to be phosphorylated in individual high throughput studies (Wang et al., 2008; Bian et al., 2014) (**Fig. 1D**). To determine whether the phosphorylation of ACBD5 at T252 (upstream of the FFAT-like motif) and S269 (within the FFAT core) affects ACBD5-VAPB interaction, we generated Myc-ACBD5 non-phosphorylatable and phosphomimetic site mutants by replacing these residues with alanine (A) or glutamic acid (E), respectively. The constructs were expressed in COS-7 cells and their interaction with endogenous VAPB was assessed by immunoprecipitation (**Fig. 3A**). Both the T252A and T252E mutant of Myc-ACBD5 bound VAPB similar to WT control levels indicating that upstream T252 phosphorylation/dephosphorylation is not crucial for ACBD5-VAPB binding. The phosphomimetic mutation of the serine in the FFAT core of ACBD5 (S269E; position 5) abolished binding to VAPB, whereas ACBD5 S269A still co-precipitated with VAPB (**Fig. 3A**).

**Figure 3.**
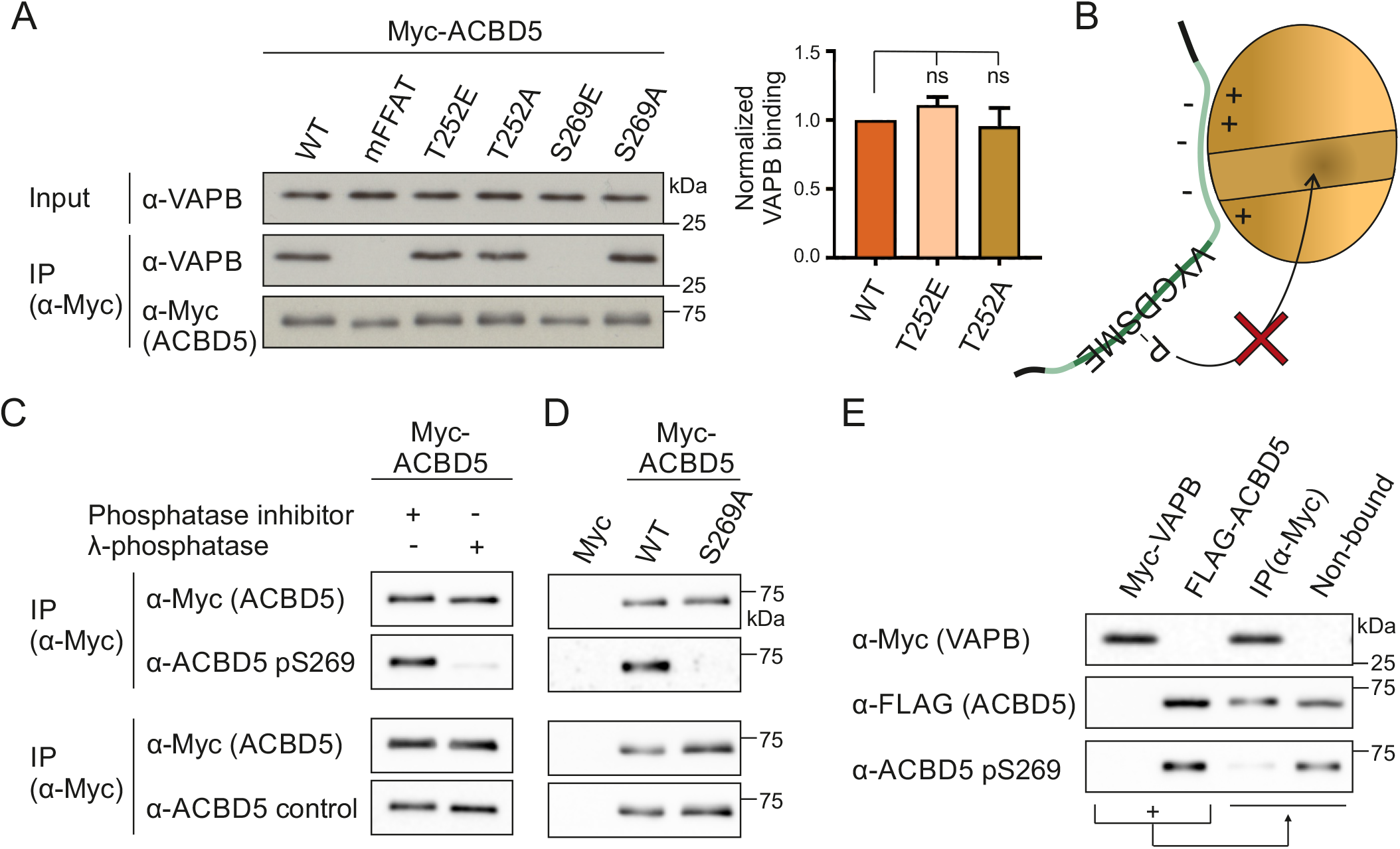
The serine in the FFAT core of ACBD5 is phosphorylated and inhibits VAPB binding. (**A**) ACBD5 constructs with non-phosphorylatable (S → A) and phosphomimetic (S → E) residues upstream (T252) or within the FFAT core (S269) were generated and expressed in COS-7 cells. The proteins were immunoprecipitated and endogenous bound VAPB detected by immunoblotting using Myc/VAPB antibodies. Data were analyzed by one-way analysis of variance with Dunnett’s multiple comparison test; ns, not significant. Error bars represent SEM. Results of three independent IPs were quantified. Total VAPB (IP fraction) was normalized against total VAPB (input) and Myc-ACBD5 (IP fraction). (**B**) The serine residue within the core of the ACBD5 FFAT-like motif (position 5, S269) binds VAPB in a hydrophobic pocket (Kaiser et al., 2005; Furuita et al., 2010). Phosphorylation at this position likely causes steric hindrance, inhibiting the FFAT-VAPB interaction. (**C**) Lysates of COS-7 cells expressing Myc-ACBD5 were treated with or without λPP before immunoprecipitation. Phosphorylation of ACBD5 at S269 was examined by immunoblotting using ACBD5 pS269/ACBD5 control/Myc antibodies. (**D**) Myc-ACBD5 WT, S269A phospho-mutant or control vector (Myc) was expressed in COS-7 cells. The proteins were immunoprecipitated and phosphorylation of ACBD5 at S269 was examined by immunoblotting using ACBD5 pS269/ACBD5 control/Myc antibodies. The ACBD5 antibodies were generated against a peptide of the same region. (**E**) Myc-VAPB and FLAG-ACBD5 were immunoprecipitated from COS-7 cells separately and subsequently incubated together to allow Myc-VAPB and FLAG-ACBD5 to interact. ACBD5 S269 phosphorylation in the VAPB-bound (IP) and non-VAPB-bound fraction was examined by immunoblotting using ACBD5 pS269/FLAG/Myc antibodies. WT, wild type; mFFAT, mutated FFAT motif (negative control).

Serine residues at similar positions in ACBD4 were investigated in parallel (**Fig. S2B**). Phosphomimetic site mutants of residues upstream of the FFAT-like motif of FLAG-ACBD4 (S166E/S169E/S171E) co-precipitated more prominently with VAPB than the FLAG-ACBD4 non-phosphorylatable mutants (S166A/S169A/S171A) when compared to WT controls (**Fig. S3B, C**). Overall, the non-phosphorylatable ACBD4 mutants appeared to have little impact on VAPB binding, whereas the ACBD4 phosphomimetic mutants increased interaction. The phosphomimetic mutant of the serine at position 5 of the ACBD4 FFAT core (S183) showed, similar to this residue in ACBD5, loss of VAPB binding (**Fig. S3B**).

### Serine 269 in the FFAT core of ACBD5 is phosphorylated

Our results indicate that phosphorylation of the serine in the FFAT core would abolish VAPB binding (**Fig. 3A, B**). To confirm that this serine in the FFAT core of ACBD5 can be phosphorylated under our experimental conditions, we generated a phospho-specific antibody towards ACBD5 pS269 (Eurogentec). As the antibody was not able to recognize ACBD5 in whole cell lysates, ACBD5 was immunoprecipitated to test the specificity of the antibody. The ACBD5 pS269 antibody showed a reduced signal in phosphatase treated lysates, while the signal of the ACBD5 control antibody (generated against a peptide of the same region), was not affected by the treatment (**Fig. 3C**). To further validate the phospho-antibody, COS-7 cells were transfected with Myc-ACBD5 WT or S269A phosphosite mutant. The ACBD5 control antibody detected both the ACBD5 WT and mutant forms, however, the pS269 antibody was not able to detect the site-specific mutant (**Fig 3D**). These experiments indicate that the generated ACBD5 pS269 antibody is specific and that the serine in the FFAT core of ACBD5 can be phosphorylated in COS-7 cells.

We now used the ACBD5 pS269 antibody to assess whether phosphorylation of the serine at position 5 of the ACBD5 FFAT core (S269) would inhibit VAPB binding. We hypothesised that only a certain fraction of ACBD5 would be phosphorylated at S269, that this population would not interact with VAPB, and thus, in an ACBD5-VAPB interaction assay the phosphorylated form would be enriched in the non-bound fraction (**Fig. 3E**). To investigate this using our pS269 antibody (which cannot detect ACBD5 in whole cell lysates), we expressed Myc-VAPB and FLAG-ACBD5 separately in COS-7 cells. Both proteins were immunoprecipitated, and subsequently incubated together to allow Myc-VAPB and FLAG-ACBD5 to interact. We then compared the phosphorylation state of FLAG-ACBD5 bound to Myc-VAPB with the non-bound fraction. Incubation with a FLAG antibody revealed approximately equal amounts of FLAG-ACBD5 in both the bound and non-bound fraction (**Fig. 3E**). However, the S269 phosphorylated form of ACBD5 was barely detectable in the VAPB-bound fraction, and was instead enriched in the non-bound fraction. This indicates that phosphorylation of ACBD5 at S269 in the FFAT core inhibits the interaction with VAPB.

### Phosphosites within the ACBD5 FFAT-like motif influence peroxisome-ER contacts

We next investigated if phosphorylation of the ACBD5 FFAT-like motif, which modulates ACBD5-VAPB binding, could also alter peroxisome-ER interactions in mammalian cells. We expressed a set of Myc-ACBD5 phosphosite mutants in ACBD5 knock-out (KO) HeLa cells and quantified peroxisome-ER association by transmission electron microscopy using unbiased spatial stereology as previously described (Costello et al., 2017b) (**Fig. 4A–C**). We have recently shown that ACBD5 KO HeLa cells have reduced peroxisome-ER associations (Bishop et al., 2019). Restoration of peroxisome-ER contacts upon ACBD5 expression was quantified by determining the mean population of peroxisomes in close contact (<15 nm) with the ER (**Fig. 4B**, mean attachment) and the proportion of the peroxisome surface closely apposed (<15 nm) to the ER (**Fig. 4C**, mean ER contact) (Costello et al., 2017b). We observed that expression of Myc-ACBD5 WT restored peroxisome-ER associations in ACBD5 KO cells to a level comparable to control HeLa cells (Bishop et al., 2019) (**Fig. 4A–C**). The S259A mutant, which did not affect ACBD5-VAPB interaction (**Fig. 2C**), restored peroxisome-ER contacts similar to ACBD5 WT, whereas expression of mutants which significantly reduced or abolished VAPB binding (non-phosphorylatable S261A and S259A/S261A/S263A, phosphomimetic S269E) (**Fig. 2C, 3A**) did not restore peroxisome-ER associations (**Fig. 4A–C**). All constructs were expressed equally well in ACBD5 KO cells (**Fig. 4D**). We have shown previously that the ACBD5-VAPB interaction plays a role in peroxisomal membrane expansion, which is a prerequisite for peroxisome division and multiplication (Schrader et al., 2016; Costello et al., 2017b). Peroxisome-ER tethering likely allows transfer of membrane lipids to promote peroxisome elongation (Schrader et al., 2020). In line with this, expression of ACBD5 WT in COS-7 cells induced the formation of tubular peroxisomes (**Fig. S3A, D**) (Hua et al., 2017). Interestingly, expression of ACBD5 S269E, which inhibits VAPB interaction and thus, peroxisome-ER contacts, did not promote peroxisome elongation. These observations further support the notion that ACBD5-VAPB mediated peroxisome-ER contacts support membrane lipid transfer by a yet unknown mechanism. Overall, this demonstrates that alterations in phosphorylated residues of ACBD5, which affect VAPB binding, also alter peroxisome-ER associations and peroxisome membrane dynamics.

**Figure 4.**
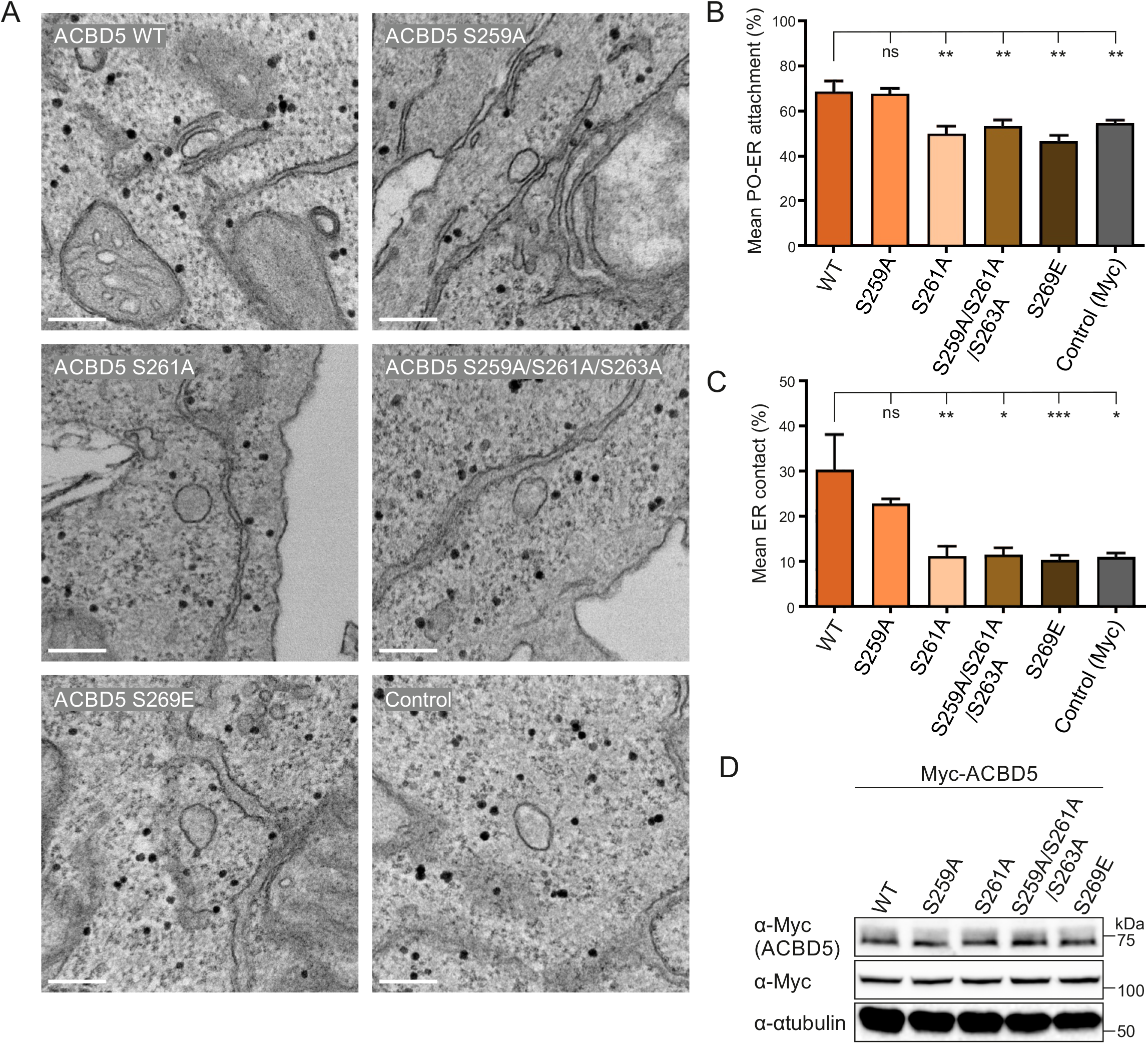
ACBD5 phospho-sites alter peroxisome-ER associations. (**A**) Representative electron micrographs of peroxisome-ER contacts in ACBD5 KO HeLa cells transfected with Myc-ACBD5 WT, S259A, S261A, S259A/S261A/S263A, S269E or control vector (Myc). (**B**) Quantitative analysis of the mean population of peroxisomes associated with the ER. (**C**) Assessment of the mean peroxisome membrane surface in direct contact with the ER. Data were analyzed by one-way analysis of variance with Dunnett’s multiple comparison test; ns, not significant; *, P < 0.05; **, P < 0.01; ***, P < 0.001. Error bars represent SEM. Results of four grids per condition. (**D**) Immunoblots of cell lysates from ACBD5 KO HeLa cells expressing the indicated Myc-ACBD5 constructs. αTubulin and Myc (unspecific band) served as loading control. Bars: 200 nm. WT, wild type.

### GSK3β alters the ACBD5-VAPB interaction

To identify kinases/phosphatases involved in the phosphorylation/dephosphorylation of ACBD5 and thus, in regulating the ACBD5-VAPB interaction, we took a candidate-based approach with focus on known associations with ACBD5 or VAPB, and previous links to peroxisome function. This approach identified glycogen synthase kinase-3 beta (GSK3β), which has recently been linked to regulation of peroxisome number in a *Drosophila* screening approach (Graves et al., 2020). GSK3β has also been linked to the regulation of PTPIP51-VAPB interaction, which forms a mitochondria-ER tethering complex (Stoica et al., 2014, 2016). To investigate a potential role for GSK3β in regulating the ACBD5-VAPB interaction, we co-expressed GSK3β and Myc-VAPB in COS-7 cells and determined alterations in the interaction of Myc-VAPB with endogenous ACBD5 and PTPIP51 (**Fig. 5A**). We confirmed that GSK3β expression increased GSK3β’s activity and downstream signalling events (e.g. β-catenin phosphorylation and degradation) (**Fig. S4A**), and reduced the VAPB-PTPIP51 interaction as previously shown (Stoica et al., 2014) (**Fig. 5A**). The interaction between ACBD5 and VAPB was also significantly reduced, suggesting that GSK3β expression altered the interaction.

**Figure 5.**
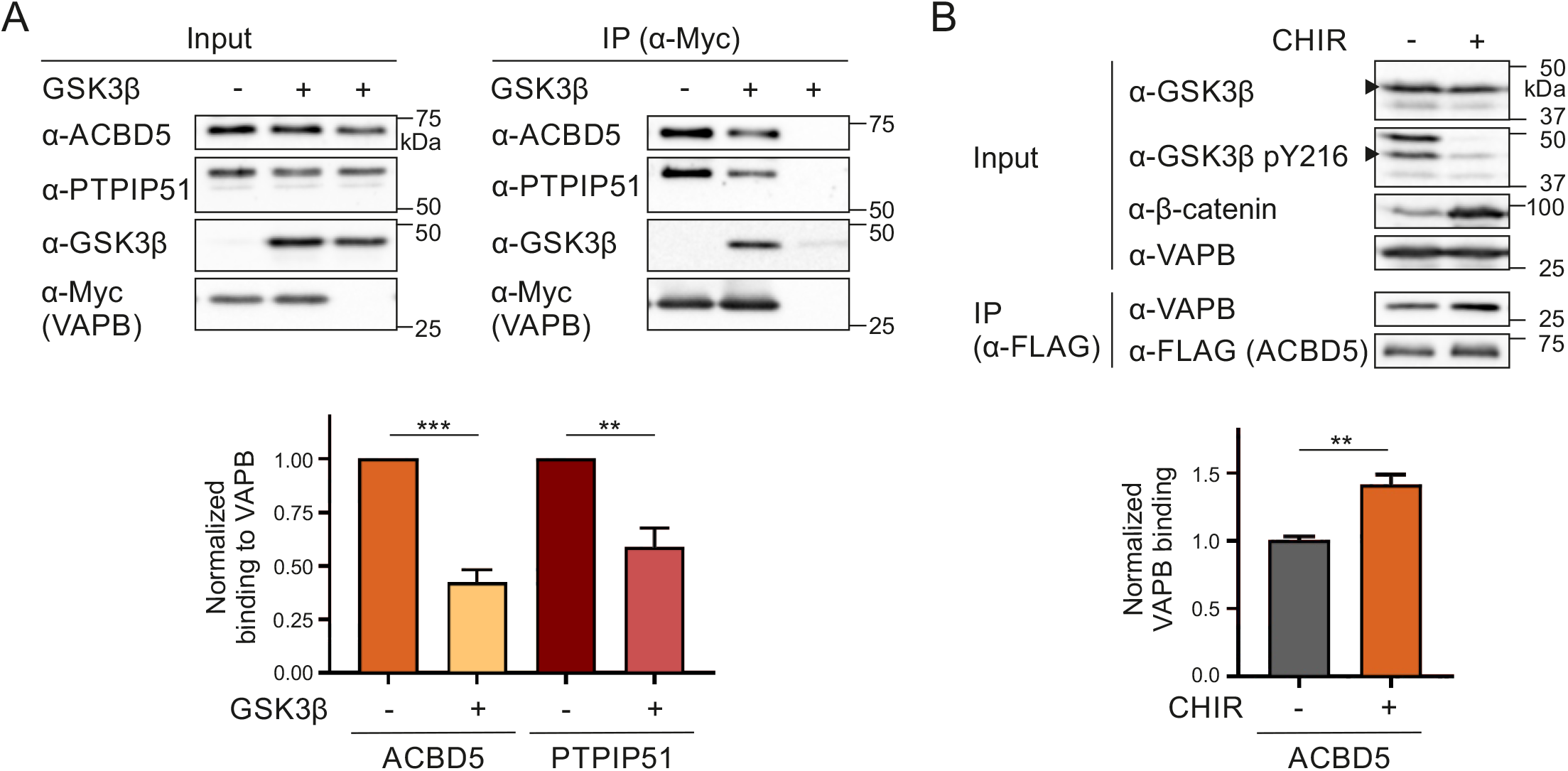
GSK3β affects the ACBD5-VAPB interaction. (**A**) Myc-VAPB (or Myc control vector) was expressed in the absence or presence of GSK3β in COS-7 cells. Myc-VAPB was immunoprecipitated and endogenous bound ACBD5 and PTPIP51 detected by immunoblotting using Myc/ACBD5/PTPIP51 antibodies. Results of three independent IPs were quantified. ACBD5/PTPIP51 (IP fraction) was normalized against total ACBD5/PTPIP51 (input) and Myc-VAPB (IP fraction). (**B**) FLAG-ACBD5 was expressed in HEK293T cells. Cells were treated with 10 μM CHIR (GSK3β inhibitor) or DMSO (control) for 16 h. FLAG-ACBD5 was immunoprecipitated and endogenous bound VAPB detected by immunoblotting using FLAG/VAPB antibodies. Inhibition of GSK3β by CHIR was confirmed using GSK3β/GSK3β pY216/β-catenin antibodies (arrow indicates GSK3β). VAPB (IP fraction) was normalized against total VAPB (input) and FLAG-ACBD5 (IP fraction). n = 5-8 of three independent IPs. Data were analyzed by a two-tailed unpaired t-test; **, P < 0.01; ***, P < 0.001. Error bars represent SEM.

To further elucidate the role of GSK3β in regulating the ACBD5-VAPB interaction, we treated HEK293T cells expressing FLAG-ACBD5 with the GSK3β inhibitor CHIR99021 (CHIR). Inhibition of GSK3β by CHIR was confirmed by decreased GSK3β Y216 phosphorylation and stabilization of its substrate β-catenin (**Fig. 5B**). Addition of CHIR significantly increased the interaction between FLAG-ACBD5 and endogenous VAPB, further indicating a role for GSK3β in regulating ACBD5-VAPB binding.

### GSK3β associates with ACBD5 and VAPB

To test for interaction of GSK3β with the ACBD5-VAPB complex, we immunoprecipitated VAPB and identified GSK3β as a potential binding partner (**Fig. 5A**), suggesting that GSK3β is present in a complex with VAPB and potentially ACBD5. We checked the protein sequence of GSK3β for potential FFAT motifs and discovered a non-canonical FFAT motif (_233_^1^DYTSSID^7^_239_) (Murphy and Levine, 2016). Although the predicted FFAT motif in GSK3β has a relatively low FFAT score (3.5; similar to ACBD4) and is located in a structured region, we generated a FFAT mutant (S237E ‘mFFAT’) which should abolish potential VAPB interaction. Expression of GSK3β S237E together with FLAG-VAPB showed that the co-immunoprecipitation of GSK3β with VAPB did not depend on the potential FFAT motif (**Fig. 6A**). To explore if GSK3β also associates with ACBD5, and if this interaction depends on VAPB, we co-expressed GSK3β and FLAG-ACBD5 WT, mFFAT or ΔTMD, a mutant with cytosolic localisation (**Fig. S3A**) in COS-7 cells. GSK3β was immunoprecipitated with all FLAG-ACBD5 variants (**Fig. 6A**), indicating that the interaction of GSK3β with ACBD5 does not depend on ACBD5-VAPB binding, the presence of ACBD5 at ER contact sites, or ACBD5 anchorage at the peroxisomal membrane. Next, we assessed if the VAPB-GSK3β co-immunoprecipitation was dependent on VAPB’s ability to bind to FFAT motif-containing proteins such as ACBD5. We observed that a FLAG-VAPB mutant unable to bind FFAT-motifs (K87D/M89D mMSP (Kaiser et al., 2005); **Fig. 6B, Fig. S4B**) still immunopreciptated with GSK3β (**Fig. 6B**). Overall, these experiments show that both ACBD5 and VAPB immunoprecipitate GSK3β, independent of their ability to interact with each other.

**Figure 6.**
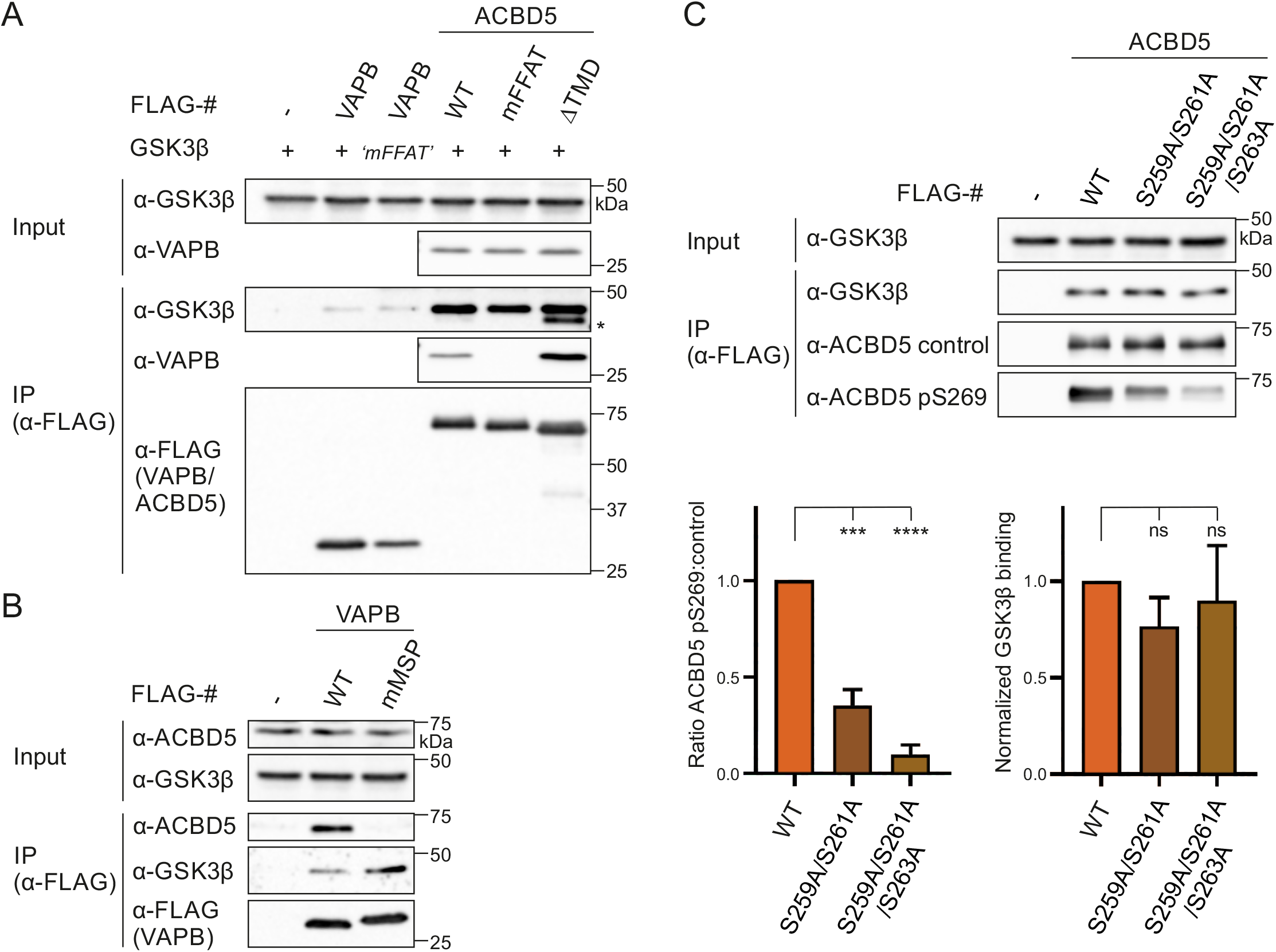
GSK3β associates with ACBD5 and VAPB. (**A**) GSK3β (S237E ‘mFFAT’) was co-expressed with FLAG-VAPB, FLAG-ACBD5 WT/mFFAT/ΔTMD, or control vector (FLAG) in COS-7 cells. FLAG-VAPB/ACBD5 were immunoprecipitated and bound GSK3β and endogenous VAPB detected by immunoblotting using FLAG/GSK3β/VAPB antibodies. Asterisk indicates unspecific band (due to reprobing of the blot). (**B**) GSK3β was co-expressed with FLAG-VAPB (K87D/M89D mMSP), or control vector (FLAG) in COS-7 cells. FLAG-VAPB w immunoprecipitated and bound GSK3β and endogenous ACBD5 detected by immunoblotting using FLAG/GSK3β/ACBD5 antibodies. (**C**) FLAG-ACBD5 constructs with non-phosphorylatable (S → A) residues in the acidic tract were co-expressed with GSK3β in COS-7 cells. FLAG-ACBD5 was immunoprecipitated and phosphorylation at S269 (pS269) was detected by immunoblotting using ACBD5 pS269 and ACBD5 control antibodies (generated against a peptide of the same region). Bound GSK3β was detected by immunoblotting using ACBD5 control/GSK3β antibodies. GSK3β (IP fraction) was normalized against total GSK3β (input) and ACBD5 control (IP fraction). Data were analyzed by one-way analysis of variance with Dunnett’s multiple comparison test; ns, not significant; ***, P < 0.001; ****, P < 0.0001. Error bars represent SEM. Results of three independent IPs were quantified. WT, wild type; mFFAT, mutated FFAT motif; TMD, transmembrane domain.

### Non-phosphorylatable residues in the acidic tract reduce ACBD5 S269 phosphorylation

A dependence of the FFAT-VAP affinity on a combination of (non)phosphorylated residues/regions in and outside the FFAT region has been suggested, which involves ‘crosstalk’ of those sites in the regulation of protein interaction and function (Goto et al., 2012; Kumagai et al., 2014). Furthermore, (de)phosphorylated regions could also be binding sites for phosphatases and kinases that act on residues further up/downstream. Therefore, we decided to assess the phosphorylation status of the serine in the FFAT core (S269) of ACBD5 in the presence of non-phosphorylatable residues in the FFAT acidic tract (**Fig. 6C**). As the acidic tract of ACBD5 resembles the consensus sequence of GSK3β (SxxxpS) (Frame and Cohen, 2001), the binding to GSK3 was also examined. Both FLAG-ACBD5 S259A/S261A and S259A/S261A/S263A showed a strong reduction in S269 phosphorylation, whereas binding to co-expressed GSK3β was not significantly altered (**Fig. 6C**). We suggest that residues in the acidic tract of ACBD5 are involved in regulating the phosphorylation of the FFAT core.

### GSK3β modulates the ACBD5-VAPB interaction via S269

As both GSK3β activity and ACBD5 S269 phosphorylation inhibit the ACBD5-VAPB interaction, we explored the possibility that GSK3β modulates the interaction via S269. We co-expressed GSK3β or a catalytically inactive mutant (GSK3β K85A) (**Fig. S4A**) with Myc-ACBD5 WT or S269A in COS-7 cells, and assessed the binding of endogenous VAPB to Myc-ACBD5 by immunoprecipitation. The binding of VAPB to Myc-ACBD5 WT was significantly reduced in the presence of GSK3β WT compared to GSK3β K85A (**Fig. 7A**). However, this reduction in VAPB binding was restored with the ACBD5 mutant S269A, suggesting that the ability of GSK3β to modulate ACBD5-VAPB interaction was dependent on the presence of a serine at position 269. To assess whether the inhibition of VAPB binding by GSK3β is linked to the phosphorylation of this serine, we next analysed the levels of ACBD5 pS269. Expression of GSK3β WT significantly increased the pS269 levels, suggesting that GSK3β inhibits the ACBD5-VAPB interaction by inducing phosphorylation at S269 of ACBD5 (**Fig. 7A**). To examine whether GSK3β potentially phosphorylates S269 directly, we developed an *in vitro* kinase assay using recombinant protein. The level of S269 phosphorylation of recombinant ACBD5 was determined in the absence and presence of recombinant GSK3β. The pS269 antibody signal was highly increased in the presence of GSK3β, showing that GSK3β can phosphorylate ACBD5 at this serine residue (**Fig. 7B**).

**Figure 7.**
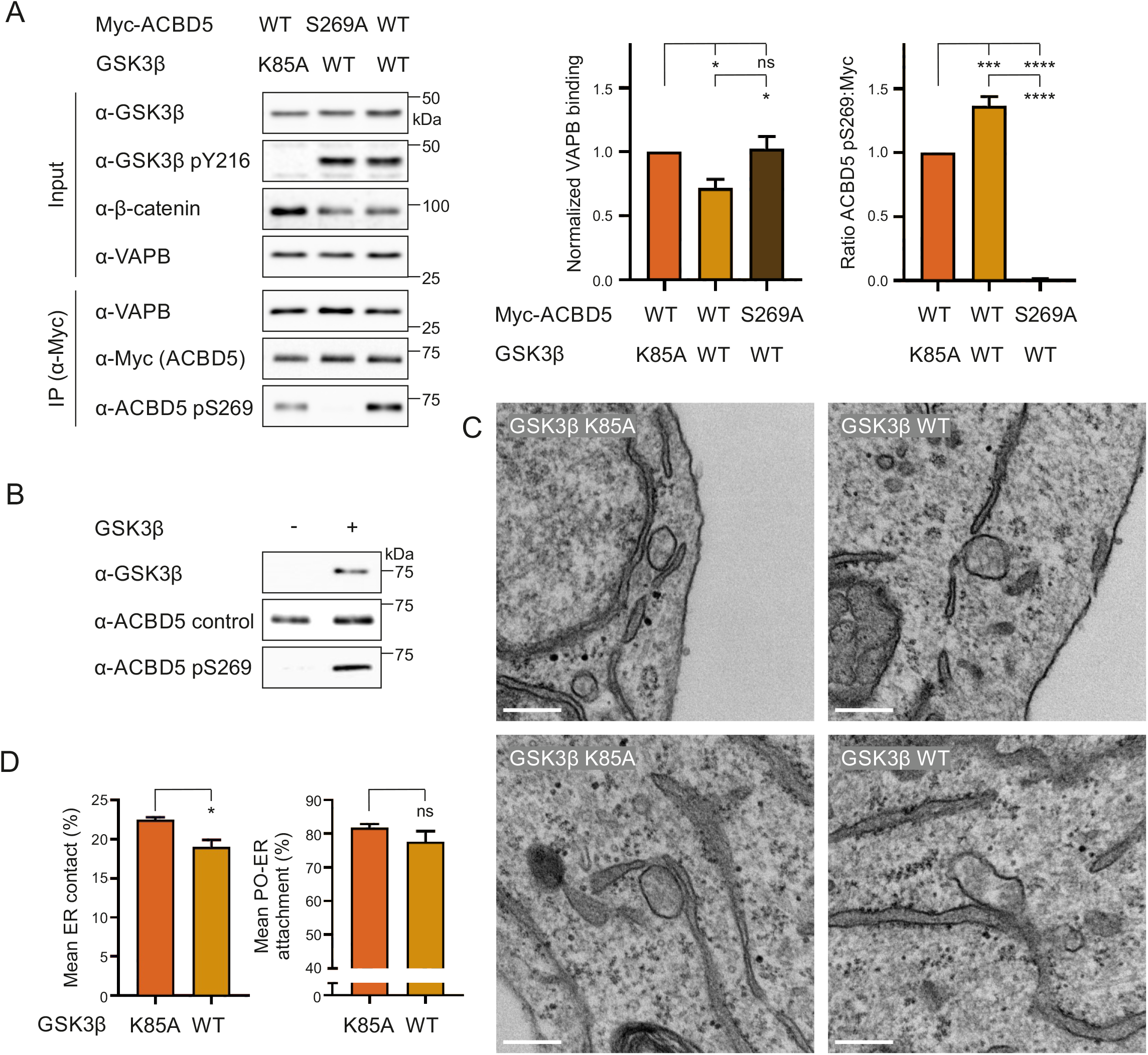
GSK3β modulates the ACBD5-VAPB interaction via S269. (**A**) GSK3β (K85A) was co-expressed with Myc-ACBD5 WT or S269A in COS-7 cells. Myc-ACBD5 was immunoprecipitated and endogenous bound VAPB detected by immunoblotting using Myc/VAPB antibodies. VAPB (IP fraction) was normalized against total VAPB (input) and Myc-ACBD5 (IP fraction). Phosphorylation of ACBD5 S269 (pS269) was detected by immunoblotting using ACBD5 pS269/Myc antibodies. GSK3β catalytic (in)activity was confirmed using GSK3β/GSK3β pY216/β-catenin antibodies. Data were analyzed by one-way analysis of variance with Tukey’s multiple comparisons test. Results of four independent IPs were quantified. (**B**) Recombinant His-ACBD5 was incubated in the absence or presence of recombinant GST-GSK3β. Phosphorylation of ACBD5 at S269 was examined by immunoblotting using ACBD5 pS269/ACBD5 control antibodies. (**C**) Representative electron micrographs of peroxisome-ER contacts in COS-7 cells transfected with a catalytically inactive GSK3β (GSK3β K85A) or GSK3β WT. (**D**) Assessment of the mean peroxisome membrane surface in direct contact with the ER. Quantitative analysis of the mean population of peroxisomes associated with the ER. Data were analyzed by a two-tailed unpaired t-test. Results of four grids per condition. ns, not significant; *, P < 0.05; ***, P < 0.001; ****, P < 0.0001. Error bars represent SEM. Bars: 200 nm. WT, wild type.

To explore if the inhibition of the ACBD5-VAPB interaction by GSK3β also alters peroxisome-ER membrane contacts, we expressed GSK3β and a catalytically inactive mutant (GSK3β K85A) in COS-7 cells and quantified peroxisome-ER association by transmission electron microscopy as described above (see **Fig. 4**). The proportion of peroxisome surface in contact with the ER was significantly reduced upon GSK3β WT expression (**Fig. 7C, D**, mean ER contact). Additionally, the number of peroxisomes in close contact with the ER also showed a slight, although not significant, reduction (**Fig. 7D**, mean attachment). We have previously shown that reduced ACBD5-VAPB interactions impact peroxisome membrane elongation in cells with impaired peroxisome division, likely because of reduced membrane lipid transport from the ER to peroxisomes (Costello et al., 2017b; Hua et al., 2017). To further investigate a role for GSK3β in inhibiting peroxisome-ER contacts, we expressed GSK3β in patient fibroblasts deficient in division factor MFF, which show highly elongated peroxisomes (Koch et al., 2016) (**Fig. S4C**). GSK3β significantly decreased the formation of these highly elongated peroxisomes, suggesting a change in peroxisome-ER membrane contacts (**Fig. S4C**). Together, these results implicate that GSK3β inhibits the ACBD5-VAPB interaction and reduces peroxisome-ER associations, required for peroxisomal membrane growth.

## DISCUSSION

We show here that peroxisome-ER association via the ACBD5-VAPB tether is regulated by phosphorylation. Several lines of evidence support this: (a) ACBD5-VAPB interaction is phosphatase-sensitive; (b) ACBD5 phosphomimetic and non-phosphorylatable mutants alter the interaction with VAPB; (c) ACBD5 phosphosite mutants impact peroxisome-ER interaction in mammalian cells; (d) GSK3β regulates the ACBD5-VAPB interaction, hence peroxisome-ER contacts; and (e) ACBD5 can be phosphorylated by GSK3β. In conclusion, our findings reveal for the first time a molecular mechanism for the regulation of peroxisome-ER contacts in mammalian cells (**Fig. 8**).

**Figure 8.**
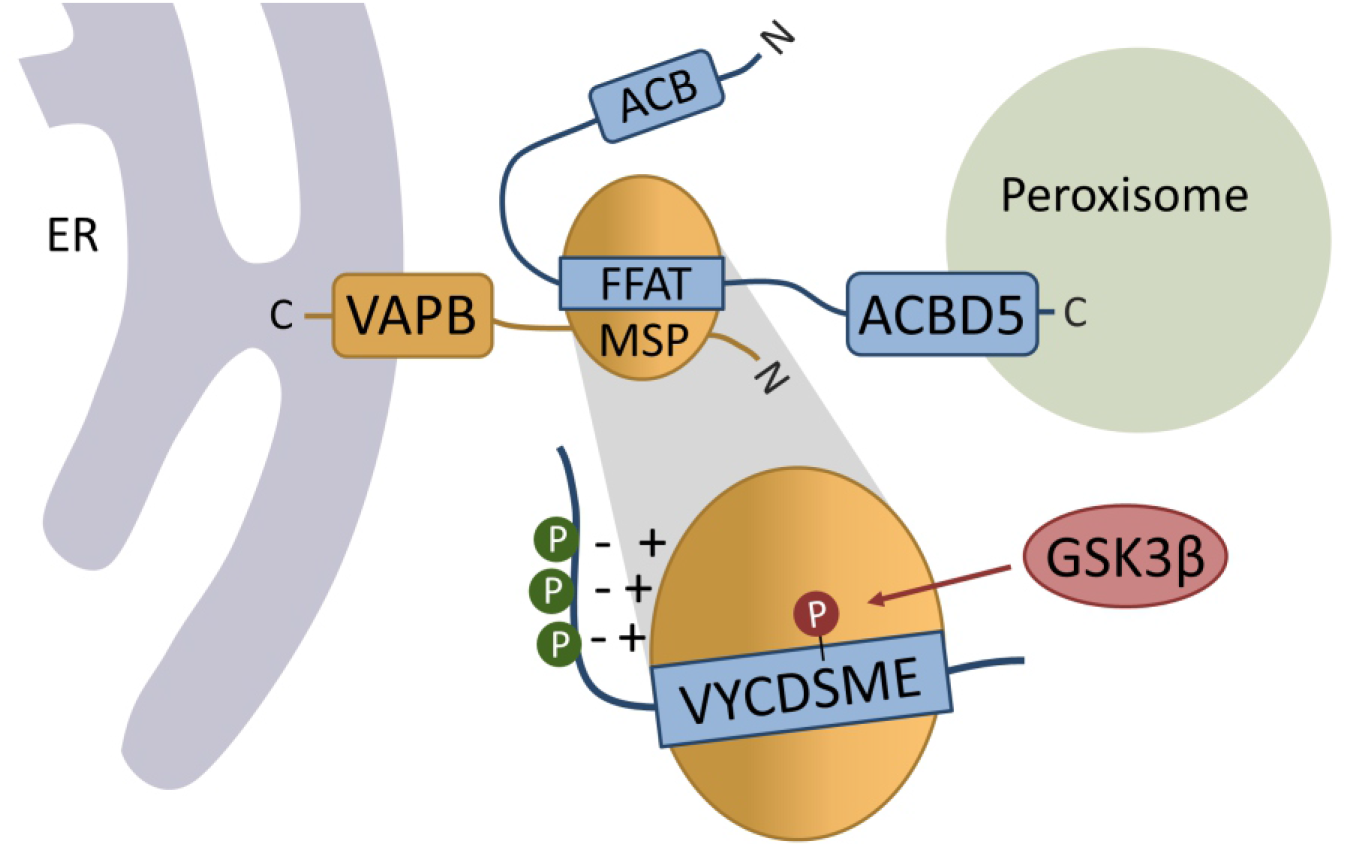
Model of ACBD5-VAPB interaction and regulation at the peroxisome-ER interface. Peroxisomal ACBD5 and ER-resident VAPB interact via the FFAT and MSP domains to enable peroxisome-ER contacts. The FFAT-MSP interaction involves the FFAT core (VYCD<u>S</u>ME) and flanking acidic tract. ACBD5 phosphorylation (P) can promote (**green; acidic tract**) or inhibit (**red; FFAT core**) VAPB interaction. GSK3β regulates the ACBD5-VAPB interaction via phosphorylation of the serine in the FFAT core. ACB, acyl-CoA binding domain.

Our findings also expand the current view and molecular understanding of FFAT motifs in general. We demonstrate that, as suggested in the original study describing FFAT motifs (Loewen et al., 2003), residues upstream of the canonical FFAT core are relevant for FFAT-VAP interactions. We show that phosphorylation of serine/threonine residues within the acidic tract (e.g. ACBD5 S261; **Fig. 2C–E**) or further extension of the acidic tract by upstream serine/threonine residues (e.g. ACBD4; **Fig. S3B, C**) improves binding to VAPB. The interaction of the FFAT motif with VAPB is thought to occur in two steps: an initial electrostatic interaction between the acidic tract upstream of the FFAT core and the basic electropositive face of the MSP domain of VAPB, followed by the binding of the FFAT core region to specific residues of the MSP domain (Furuita et al., 2010) (**Fig. 2B; Fig. 8**). Our findings suggest that the FFAT motif can be “activated” by adding a negatively charged phosphate group to serine/threonine residues, improving binding of the acidic tract to the basic electropositive face of the MSP domain. Our data on ACBD4 phosphorylation suggest that these residues can also potentially be located further upstream, thus extending the canonical FFAT-like motif, as the ACBD4 phosphomimetic mutants increased ACBD4-VAPB binding (**Fig. S3B, C**). These residues have been found to be phosphorylated in high-throughput studies (Hornbeck et al., 2015). However, in our experimental system non-phosphorylatable mutants (S166A/S169A/S171A) (**Fig. S3B, C**) and phosphatase treatment (**Fig. 1B, Fig. S1A**) did not have a significant impact on ACBD4-VAPB binding, suggesting low phosphorylation levels in our experimental setup. We therefore propose that the extended site may function as a switch to modulate ACBD4-VAPB binding potentially dependent on particular cellular conditions.

Numerous FFAT motif-containing proteins possess serine/threonine residues in their acidic tract, including ceramide transport protein CERT, lipid transfer protein STARD3 and potassium channel Kv2. Phosphorylation of a single serine residue (S315) in the acidic tract of CERT has been reported to enhance VAPA interaction (Kumagai et al., 2014). However, mutation of this residue to S315A did not have much impact on VAPA interaction. The interaction between STARD3 FFAT peptide and endogenous VAPA/B was strengthened by phosphorylation of a serine residue in both the upstream and downstream flank of the acidic tract (Di Mattia et al., 2020). Kv2 contains a FFAT motif that without phosphorylation would not have an ‘acidic tract’ (Kirmiz et al., 2018; Johnson et al., 2018). Another example of such a protein is *Chlamydia trachomatis* IncV which contains ‘acidic tracts’ exclusively composed of serine residues (Stanhope et al., 2017). This suggests that overall phosphorylation is a general mechanism to “activate” the acidic tract and by doing so adjust VAP-interactions. As there are a large number of VAP interactors, and evidence of competition between FFAT proteins, these small increases in VAP affinity may have a significant impact. The importance of the residues in the acidic tract can also be illustrated by the identification of the unconventional FFAT-related FFNT [two phenylalanines (FF) in a neutral tract] motifs, that preferably bind to ER-resident MOSPD1/3 proteins that, like VAPA/B, possess a MSP domain (Cabukusta et al., 2020). Moreover, PTPIP51 contains an ‘acidic tract’ mainly composed of serine/threonine residues and was able to bind all VAP and MOSPD proteins. (De)phosphorylation of the residues in the tract could possibly switch the tract from acidic to neutral and *vice versa*, changing the affinity of PTPIP51 for VAPA/B and MOSPD1/3. However, our data also imply that there are potentially additional phosphorylation events involved in the regulation of the ACBD5-VAPB interaction. Phosphorylation of VAPB may also influence ACBD5-VAPB interaction (Bian et al., 2014; Sharma et al., 2014). Furthermore, dimerisation has been implicated to enhance FFAT-VAPB interaction (Kaiser et al., 2005; Mikitova and Levine, 2012).

In summary, we suggest that the negatively charged phosphate groups support the overall acidic environment by increasing affinity for the electropositive face of the VAP-MSP domain, thus facilitating FFAT-MSP interaction (**Fig. 2B, Fig. 8**). An alternative explanation may be that phosphorylation induces structural rather than electrostatic changes, which enhance VAP binding. An extended acidic tract may be particularly important for VAP binding of proteins with “weaker” FFAT motifs. The prediction of non-canonical FFAT motifs is problematic [e.g. the ACBD4 FFAT-like motif has a FFAT score of 3.5, which is outside the generally-used cut-off of 2.5 (Murphy and Levine, 2016; Slee and Levine, 2019)]. It is possible that serine/threonine residues within and upstream of the acidic tract contribute more prominently to VAP affinity than is currently factored into the scoring algorithm (Murphy and Levine, 2016). Such residues may therefore need to be considered so that proteins with a lower FFAT score are not omitted.

Substitution of the serine residue at position 5 of the FFAT core to phosphomimetic glutamic acid in both ACBD4 and ACBD5 abolished their binding to VAPB completely (**Fig. S3B, Fig. 3A**). A previous study, in which peptides of human proteins were expressed in yeast, reported that glutamic acid at this position in the FFAT core of human AKAP220 completely inhibited ER-localisation, suggesting a loss of binding to the yeast VAP homolog Scs2p (Mikitova and Levine, 2012). Furthermore, we show that the serine at position 5 of the ACBD5 FFAT core (S269) can be phosphorylated (**Fig. 3C, D**) and that this phosphorylated population is only detectable in the fraction not immunoprecipitated by VAPB, indicating that FFAT phosphorylation at position 5 blocks VAP interaction. (**Fig. 3E**). The canonical FFAT motif has an alanine residue at this position (^1^EFFDA-E^7^), which binds VAP in a small hydrophobic pocket (Furuita et al., 2010; Kaiser et al., 2005) (**Fig. 2B**). Steric hindrance likely excludes glutamic acid and phosphorylated serine from this pocket, and thereby binding of the peptide/protein to VAP. At first glance, phosphorylation within the FFAT-like motif that inhibits VAPB binding appears to be in contrast to our observation that phosphatase treatment/dephosphorylation of ACBD5 inhibits VAPB binding (see **Fig. 1**). However, phosphorylation of residues within the acidic tract (outside the core region) is likely required to initiate a more transient ACBD5-VAPB interaction prior to interaction of the FFAT core with the MSP hydrophobic pocket (**Fig. 2**). We speculate that phosphorylation of the serine residue within the FFAT core might represent an additional regulatory mechanism to modulate ACBD5-VAPB interaction. Importantly, phosphorylation of the serine residue at position 5 may also negatively regulate VAP interaction of other proteins with a FFAT motif; numerous FFAT motif-containing proteins possess a serine or threonine at this position (**Fig. S5**) (Murphy and Levine, 2016).

Phosphorylation upstream of the FFAT motif may also alter the phosphorylation status of the critical serine residue in the core of the ACBD4/5 FFAT-like motif (position 5). ACBD5 with non-phosphorylatable residues in the acidic tract showed reduced levels of pS269 (**Fig. 6C**). ‘Crosstalk’ between (non)phosphorylated residues/regions, in and outside the FFAT region in the regulation of the FFAT-VAP interaction and protein function has been suggested previously (Goto et al., 2012; Kumagai et al., 2014).

Interestingly, recent studies revealed that phosphorylation of serine and threonine residues at position 4 of the FFAT core strongly increases the affinity of unconventional motifs to VAP (Di Mattia et al., 2020; Guillén-Samander et al., 2021), showing that phosphorylation of the core can affect the FFAT-VAP binding in paradoxical ways, dependent on the position of the residue. The canonical FFAT motif possesses aspartic acid (D) at position 4, which resembles phosphorylated serine/threonine to enable VAP interaction, while the canonical FFAT motif possesses alanine (A) at position 5, which when replaced by phosphorylated serine/threonine inhibits VAP interaction. SNX2 and CALCOCO1, two confirmed VAP interactors (Dong et al., 2016; Nthiga et al., 2020), have serine/threonine residues at both position 4 and 5 of their FFAT motif (**Fig. S5**), suggesting that VAP binding of these proteins could be tightly regulated by two opposing phosphorylation mechanisms.

We demonstrate that GSK3β is in a complex with ACBD5 and VAPB (**Fig. 6**) and that the kinase regulates the binding of ACBD5 to VAPB; expression of GSK3β decreased the ACBD5-VAPB interaction, whereas inhibition of GSK3β increased it (**Fig. 5**). Inhibition of the ACBD5-VAPB binding by GSK3β is dependent on the serine in the FFAT core of ACBD5 (S269) (**Fig. 7A**). Furthermore, we show that GSK3β can directly phosphorylate ACBD5 at this residue *in vitro* (**Fig. 7B**). Hence, we show that expression of GSK3β alters peroxisome-ER associations (**Fig. 7C, D, Fig. S4C**). Overall, our data indicates that GSK3β negatively regulates the ACBD5-VAPB interaction by phosphorylating ACBD5 at S269. However, GSK3β acts on a large number of substrates, and its role in ACBD5-VAPB regulation could be via multiple levels (Frame and Cohen, 2001). Phosphorylation of VAPB was increased upon AKT inhibition, an upstream regulator of GSK3β (Wiechmann et al., 2021). Other protein kinases and phosphatases directly acting on ACBD5 await identification.

The previous studies showing that GSK3β activity regulates the PTPIP51-VAPB interaction, and thus mitochondria-ER associations, also linked the regulation of GSK3β activity to TDP-43 and FUS, two proteins associated with amyotrophic lateral sclerosis (ALS) and fronto-temporal dementia (FTD) (Stoica et al., 2014, 2016). This suggests that alterations of both mitochondrial-ER and peroxisome-ER contacts might be a feature of TDP-43/FUS-induced pathology. Recently, lipid alterations in the frontal cortex of patients with ALS/FTD-TDP-43 proteinopathy have been related to peroxisome impairment (Andrés-Benito et al., 2021). Several PE- and PC-plasmalogens were found to be down-regulated, similar to decreased levels reported in ACBD5 deficient patient cell lines and ACBD5 depleted cells (Hua et al., 2017; Herzog et al., 2018). As plasmalogen biosynthesis, which is initiated in peroxisomes and completed in the ER, requires peroxisome-ER cooperation, altered peroxisome-ER contacts could contribute to TDP-43-induced pathology. Aberrant activity of GSK3β has also been linked to peroxisomal disorders; GSK3β activity was increased in the nervous system of mouse models for adrenoleukodystrophy (ALD) and rhizomelic chondrodysplasia punctata (RCDP) and in patient fibroblasts (da Silva et al., 2014; Ranea-Robles et al., 2018). Whether peroxisome-ER contacts are impaired under those conditions awaits further research.

Peroxisome-ER contact sites mediated by ACBD5-VAPB have been implicated in the regulation of peroxisome motility and positioning, membrane expansion and biogenesis of peroxisomes, as well as metabolic cooperation (e.g. in plasmalogen synthesis) (Costello et al., 2017b; Hua et al., 2017). Switching peroxisome-ER tethering “ON” and “OFF” would allow regulation of these processes under different physiological conditions. For example, the control of peroxisome motility and positioning may be critical for cell division, in which peroxisome inheritance plays a role in normal cell mitosis and differentiation (Asare et al., 2017). The physiological function(s) of ACBD4-VAPB mediated peroxisome-ER contacts remain to be elucidated; they may differ from those of ACBD5-VAPB, as ACBD4 responds differently to (de)phosphorylation.

Many peroxisomal proteins, involved in various processes such as peroxisome biogenesis and protein import, are phosphorylated according to phospho-proteomics studies, but the biological function of this phosphorylation remains largely unknown (Oeljeklaus et al., 2016). Phosphorylation of human PEX5, the shuttling import receptor for peroxisomal matrix proteins, has been shown to be implicated in pexophagy in response to ROS (Zhang et al., 2015). Phosphorylation of mammalian PEX14, a membrane protein that is part of the peroxisomal import machinery, suppressed the import of catalase into peroxisomes under oxidative stress conditions and in mitotic cells (Okumoto et al., 2020; Yamashita et al., 2020). Additionally, Pex14p phosphorylation in the yeast *S. cerevisiae* controls the import of Cit2p, the peroxisomal isoform of citrate synthase (Schummer et al., 2020). Other studies in yeast also provide insight into the physiological role of phosphorylation of peroxisomal proteins. In *S. cerevisiae* and *P. pastoris*, the peroxisome biogenesis factor Pex11p is activated by site-specific phosphorylation (Knoblach and Rachubinski, 2010; Joshi et al., 2012), although this appears to be different in *H. polymorpha* (Thomas et al., 2015). In addition, pexophagy requires the phosphorylation of pexophagy receptor Atg30 in *P. pastoris* (Burnett et al., 2015). Our novel findings on the regulation of the ACBD5-VAPB tether, and subsequent peroxisome-ER membrane contacts, represent another example for a physiological role of phosphorylation of peroxisomal membrane proteins in mammals.

## MATERIALS AND METHODS

### Plasmids and antibodies

See Table S1 for details of plasmids used in this study, Table S2 for plasmids generated in this study, Table S3 for gene synthesis of ACBD4 with ACBD5 FFAT-like motif region, Table S4 for gene synthesis of ACBD5 codon optimised for *E. coli*, and Table S5 for details of primers used. Site-directed mutagenesis to generate point mutations was done using the QuikChange Site-Directed Mutagenesis kit (Agilent Technologies). All constructs produced were confirmed by sequencing (Eurofins Genomics).

The polyclonal rabbit phospho-ACBD5 Ser269 antibody (α-ACBD5 pS269) was produced by Eurogentec (Seraing, Belgium). The antibody was raised against peptide ^264^EVYCDSMEQFGQE^276^ including a phospho-Ser269. Phospho-specific and non-phospho-specific (α-ACBD5 control) antibodies targeting the peptide were purified from serum by double affinity purification. Details on antibodies used in this study can be found in Table S6.

### Cell culture, transfection and drug treatment

COS-7 (African green monkey kidney cells, CRL-1651; ATCC), HEK293T (Human embryonic kidney 293T cells; ECACC), ACBD5 KO HeLa cells (generated by J. Koster, University of Amsterdam, Netherlands) (Ferdinandusse et al., 2017) and MFF-deficient fibroblasts (provided by F.S. Alkuraya, King Faisal Specialist Hospital and Research Center, Riyadh, Saudi Arabia) (Shamseldin et al., 2012; Costello et al., 2017b) were cultured in Dulbecco’s Modified Eagle’s Medium (DMEM), high glucose (4.5 g/L) supplemented with 10% fetal bovine serum (FBS), 100 U/ml penicillin and 100 μg/ml streptomycin (all from Life Technologies) at 37°C with 5% CO_2_ and 95% humidity. COS-7 and HEK293T cells were transfected using diethylaminoethyl (DEAE)-dextran (Sigma-Aldrich) as described (Bonekamp et al., 2010) and HeLa cells with Lipofectamine 3000 (Invitrogen). HEK293T cells were seeded in dishes coated with Collagen R solution 0.4% (1:10, Serva). Transfection of fibroblasts was performed using the Neon Transfection System (Thermo Fisher Scientific) as previously described (Costello et al., 2017b). Cells were assayed for immunofluorescence or immunoblotting and immunoprecipitation experiments 24 or 48 h after transfection, respectively. To inhibit GSK3β activity, cells were treated with 10 μM CHIR99021 (Sigma-Aldrich; 5 mM stock in dimethyl sulphoxide [DMSO]) and incubated for 16 hours before immunoprecipitation. Control cells were incubated with the same volume of DMSO.

### Immunofluorescence and microscopy of cultured cells

Cells grown on glass coverslips were fixed with 4% paraformaldehyde (PFA; in PBS, pH 7.4) for 20 min, permeabilised with 0.2% Triton X-100 for 10 min, and blocked with 1% BSA for 10 min. Blocked cells were sequentially incubated with primary and secondary antibodies (Table S6) for 1 h in a humid chamber at room temperature. Coverslips were washed with ddH2O to remove PBS and mounted on glass slides using Mowiol medium. Cell imaging was performed using an Olympus IX81 microscope equipped with an UPlanSApo 100x/1.40 oil objective (Olympus Optical). Digital images were taken with a CoolSNAP HQ2 CCD camera and adjusted for contrast and brightness using MetaMorph 7 (Molecular Devices).

### Electron microscopy and spatial stereology

Electron microscopy was performed as previously described (Costello et al., 2017b). In brief, monolayers of cells were fixed for a total of 1 h in 0.5% glutaraldehyde in 0.2M PIPES buffer (pH 7.2), and post-fixed in 1% osmium tetroxide (reduced with 1.5% w/v potassium ferrocyanide) in cacodylate buffer for 1h. After washing in deionised water the cells were dehydrated in a graded ethanol series before embedding in Durcupan resin (Sigma Aldrich). 60 nm ultra-thin sections were collected on pioloform-coated 100 mesh copper EM grids (Agar Scientific) and contrasted with lead citrate prior to imaging using a JEOL JEM 1400 transmission electron microscope operated at 120 kV. Images were taken with a digital camera (ES 1000W CCD, Gatan). Quantification of peroxisome-ER contacts was performed as previously (Costello et al., 2017b). In brief, peroxisomes were sampled (n= 36-55, mean = 44±1 (**Fig. 4)** or n = 106-116, mean = 112 ± 1.46 (**Fig. 7**) peroxisomes per experimental grid) by scanning the EM grids systematic uniform random. To estimate the mean fraction of total peroxisome membrane surface in direct contact with the ER, a stereological approach by line intersection counting was used. Intersections were classified as direct membrane contact (defined as “attachment”) if there was <15 nm distance between peroxisome and ER membranes.

### Protein extraction and phosphatase treatment

Transfected cells and controls were washed in PBS, and lysed in ice-cold lysis buffer (50-100 mM Tris-HCL pH 7.4, 150 mM NaCl, 2mM DTT, 1% Triton X-100, and mini protease inhibitor cocktail (Roche), with or without phosphatase inhibitor cocktail (Roche)). Insolubilized material was pelleted by centrifugation at 15,000x g. Clarified lysates were incubated with 0.9 mM MnCl_2_ and 7 U/μl lysate lambda protein phosphatase (λPP; New England BioLabs), or 0.9 mM MnCl_2_ and H_2_O as control, for 1 h at 30°C. The total protein concentration of the lysate was determined by a Bradford protein assay (Bio-Rad). Reactions were stopped and proteins were denatured in Laemmli buffer for 10 min at 95°C.

### SDS-PAGE and immunoblotting

Proteins were separated on 10% or 12.5% conventional SDS-PAGE gels, and subsequently transferred to nitrocellulose membranes (Amersham Bioscience) using a semi-dry apparatus (Trans-Blot SD, Bio-Rad). Phos-tag, in complex with a divalent cation, binds reversibly to phosphate groups, separating phosphorylated and non-phosphorylated forms of proteins when added to polyacrylamide gels (Kinoshita et al., 2006). For Phos-tag SDS-PAGE, 50 μM Phos-tag acrylamide (Wako Chemicals, purchased from NARD Institute ltd.) and 100 μM MnCl_2_ were added to the resolving gel solution of 8% SDS-PAGE gels before polymerization. To improve electrotransfer of ACBD5, a two-layer Phos-tag SDS-PAGE method was used (Longoni et al., 2015); the Phos-tag is only present in the upper layer of the resolving gel. For two-layer Phos-tag SDS-PAGE, the same amounts of Phos-tag and MnCl_2_ were added to 6% acrylamide resolving gel solution. One volume of the Phos-tag resolving gel solution was drawn into a serological pipet. Using the same pipet, three volumes of normal 6% acrylamide resolving gel solution was drawn up. Ejection of the gel solution between the glass plates resulted in a gel with only the Phos-tag ligand in the top of the gel. Conventional SDS-PAGE was performed in parallel, including MnCl_2_ in the corresponding layers as control. After protein separation at a constant voltage of 75 V for 3 h, the Phos-tag SDS PAGE gel was incubated in transfer buffer containing 10 mM EDTA to remove Mn^2+^. Proteins were transferred to activated PVDF membranes (GE Healthcare) using the semi-dry method for 1h at 24 V. Membranes were blocked in 5% dry milk (Marvel) in Tris-buffered saline with Tween-20 (TBS-T), and incubated with primary antibodies (Table S6), followed by incubation with horseradish peroxidase-conjugated secondary antibodies (Table S6) and detected with enhanced chemiluminescence reagents (Amersham Bioscience) using Amersham hyperfilm (GE Healthcare) or the G:Box Chemi (Syngene). For antibody reprobing, membranes were incubated 2-3 times for 10-15 min in a mild membrane stripping buffer (1.5% w/v glycine, 0.1% w/v SDS, 1% v/v Tween-20, pH 2.2 (HCl)). Following this, the membranes were washed in TBS-T and blocked in 5 % dry milk in TBST-T prior to antibody incubation.

### Immunoprecipitation (IP) for *in vitro* binding assays

For *in vitro* binding assays (**Fig. 1B**), GST-VAPB was expressed in BL21 Rosetta (DE3) cells (EMD Millipore) induced with 1 mM IPTG for 4 h. Cells were centrifuged at 5,000x g for 10 min and pellets resuspended in ice-cold *Escherichia coli* lysis buffer (50 mM Tris-HCl, pH 7.5, 300 mM NaCl, 0.1% NP-40, 1 mM PMSF, and protease inhibitor cocktail). Cells were disrupted by sonication, and insoluble material was removed by centrifugation at 15,000x g. The supernatant was incubated with GST-TRAP agarose beads (ChromoTek) for 2 h at 4°C. COS-7 cell lysates from FLAG-ACBD4/5 (mFFAT) expressing cells were treated for 1 h with λPP (New England BioLabs) as described above, and subsequently incubated with the GST-VABP-bound beads for 30 min. Beads were then washed extensively with wash buffer (50mM Tris-HCL pH 7.4, 150 mM NaCl, 2mM DTT, 1% Triton X-100) in a rotating shaker at 4°C and by centrifugation at 2,500x g. Proteins were eluted with Laemmli buffer for 10 minutes at 95°C. Immunoprecipitates and total lysates were analysed by Western immunoblotting.

### IP for other binding assays

For binding assays (**Fig. 1E, S1A**), COS-7 cell lysates from FLAG-ACBD4(wACBD5_FFAT)/5 (mFFAT) expressing cells were treated for 1 h with λPP (New England BioLabs) as described above. Subsequently, DTT and Triton X-100 concentrations were adjusted to 0.4 mM and 0.2%, respectively, by using dilution buffer (50-100 mM Tris-HCL pH 7.4, 150 mM NaCl). The samples were incubated with anti-FLAG M2 affinity gel (Sigma) at 4°C for 1 h, after which the gel was repeatedly washed with dilution buffer in a rotating shaker at 4°C and by centrifugation at 5,000x g. Proteins were competitively eluted using 3X FLAG peptide (Sigma; in 10 mM Tris HCl, 150 mM NaCl, pH7.4 (TBS)). Immunoprecipitates and total lysates were analysed by Western immunoblotting.

For quantification of ACBD5-VAPB binding (**Fig. 1C**), Myc-ACBD5 was expressed in COS-7 cells and cells lysed, compatible with λPP-treatment, as described above. Insolubilized material was removed by centrifugation at 100,000x g for 20 min at 4°C. Clarified lysates were treated for 1 h with λPP (New England BioLabs) as described above. Subsequently, DTT and Triton X-100 concentrations were adjusted to 0.66 mM and 0.33%, respectively, by using dilution buffer (50 mM Tris-HCL pH 7.4, 150 mM NaCl). Lysates were then mixed with Myc-TRAP magnetic agarose beads (ChromoTek) and incubated for 1 h at 4°C on a rotating wheel. Beads were washed extensively with dilution buffer and bound proteins were eluted with Laemmli buffer for Western immunoblotting.

### IP of phospho-mutants and GSK3β experiments

For immunoprecipitation of FLAG-ACBD4 phospho-mutants (**Fig. S3A, B**) or FLAG-ACBD5/VAPB (**Fig. 5B, Fig. 6**), the constructs mentioned in the experiments were expressed in COS-7 cells. Cells were washed in PBS, and lysed in ice-cold lysis buffer (50 mM Tris-HCl pH7.4, 150 mM NaCl, 1 mM EDTA, 1% Triton X-100, mini protease inhibitor cocktail (Roche), and phosphatase inhibitor cocktail (Roche)). Insolubilized material was pelleted by centrifugation at 15,000x g. The supernatant was incubated with anti-FLAG M2 affinity gel (Sigma) and further processed as described above (beads were washed with FLAG wash buffer: 50 mM Tris-HCl pH7.4, 150 mM NaCl, 1 mM EDTA, 1% Triton X-100).

For immunoprecipitation of Myc-ACBD5 phospho-mutants (**Fig. 2C, 3A, D, 7A, S1B**) or Myc-VAPB (**Fig. 5A, S4B**), the constructs mentioned in the experiments were expressed in COS-7 cells. After 48 h cells were washed in PBS, and lysed in ice-cold lysis buffer (10 mM Tris-HCL pH 7.4, 150 mM NaCl, 0.5 mM EDTA, 0.5% NP-40, mini protease inhibitor cocktail (Roche), and phosphatase inhibitor cocktail (Roche)). Insolubilized material was pelleted by centrifugation at 15,000x g. The supernatant was diluted (1:2) with dilution buffer (10 mM Tris-HCL pH 7.4, 150 mM NaCl, 0.5 mM EDTA) and mixed with Myc-TRAP (ChromoTek) magnetic agarose beads and incubated for 1 h at 4°C. Beads were subsequently extensively washed with dilution buffer in a rotating shaker at 4°C. Proteins were eluted with Laemmli buffer for 10 min at 95°C. Immunoprecipitates and total lysates were analysed by Western immunoblotting. For immunoprecipitation with phosphatase treatment (**Fig. 2D, E, 3C**), clarified lysates were treated for 1 h with λPP (New England BioLabs) as described above. DTT and Triton X-100 concentrations were adjusted to 0.4 mM and 0.2%, respectively, by using dilution buffer (100 mM Tris-HCL pH 7.4, 150 mM NaCl), prior to beads incubation. The samples were further processed as described above.

### IP of FLAG-ACBD5 binding to Myc-VAPB

To assess the binding of FLAG-ACBD5 pS269 to Myc-VAPB (**Fig. 3E**), Myc-VAPB and FLAG-ACBD5 were expressed separately in COS-7 cells. Myc-VAPB was immunoprecipitated as described above. Beads were extensively washed with Myc wash buffer (10 mM Tris-HCL pH 7.4, 150 mM NaCl, 0.5 mM EDTA, 0.05% NP-40). FLAG-ACBD5 was immunoprecipitated as described above. The washes with FLAG wash buffer were followed by a wash in Myc wash buffer. Proteins were competitively eluted using 3X FLAG peptide (Sigma; in Myc wash buffer). The eluted FLAG-ACBD5 was incubated with the Myc-VAPB-bound beads for 1 h at 4°C. The supernatant was transferred to a new tube and the beads were subsequently extensively washed with Myc wash buffer in a rotating shaker at 4°C. Proteins were eluted with Laemmli buffer for 10 min at 95°C. Immunoprecipitates and total lysates were analysed by Western immunoblotting.

### *In vitro* kinase assay

His-ACBD5 was expressed in BL21 Rosetta (DE3) cells (EMD Millipore) induced with 1 mM IPTG overnight at 18°C. Cells were centrifuged at 5,000x g for 5 min and pellets re-suspended in ice-cold lysis buffer (50 mM Tris-HCl, pH 7.4, 300 mM NaCl, 10 mM Imidazole, 4 mM DTT, 1% Triton X-100 and protease inhibitor cocktail (Roche)). Cells were disrupted by sonication, and insoluble material was removed by centrifugation at 13,000x g. The supernatant was incubated with HisPur™ Ni-NTA agarose beads (Thermo Fisher) for 1 h at 4°C. Beads were washed extensively in wash buffer (50 mM Tris-HCl, pH 7.4, 300 mM NaCl, 25 mM Imidazole, 4 mM DTT and protease inhibitor cocktail (Roche)) to remove unbound protein. Purified His-ACBD5 was eluted from the beads by incubating with elution buffer (50 mM Tris-HCl, pH 7.4, 300 mM NaCl, 250 mM Imidazole, 4 mM DTT) for 15 min at room temperature. His-ACBD5 concentration was adjusted to 10ng/μl by diluting in kinase reaction buffer (10 mM Tris-HCl, pH 7.4, 50 mM NaCl, 5 mM MgCl2, 0.5 mM EDTA and 0.05% NP-40). For in vitro kinase assays (**Fig. 7B**), reaction mixes were prepared using 2.5 μg recombinant His-ACBD5 with or without the addition of 0.3 mM ice-cold ATP (Thermo Fisher) and 0.1 ug GST-GSK3β (Abcam). Reactions were incubated at 37°C for 30 min. Samples were prepared with Laemmli buffer and then analysed by Western immunoblotting.

### Mass spectrometry (MS)

FLAG-ACBD5 expressed in COS-7 cells was immunoprecipitated using anti-FLAG M2 affinity gel (Sigma) as described above. Subsequently, beads were washed twice in ammonium bicarbonate (50 mM) and cysteine residues were reduced with 5 mM tris(2-carboxyethyl)phosphine (20 min, 800 rpm, 37°C) and alkylated with 50 mM 2-chloroacetamide (20 min, 800 rpm, 25°C). Proteins were digested on-bead either with sequencing-grade trypsin (1:50) (Promega, Walldorf, Germany) for 4 h at 800 rpm and 42°C or thermolysine (1:50) (Promega, Walldorf, Germany) for 2 h at 800 rpm and 60°C. Peptides were acidified using TFA at a final concentration of 1% and phosphopeptide enrichment was performed using titanium dioxide (TiO2) as described previously with slight modifications (Humphrey et al., 2018). Before incubation with proteolytic peptide samples, TiO2 beads were washed using elution and wash buffer. C8 stage tips were pre-equilibrated with methanol and wash buffer. For mass spectrometric analysis, enriched and non-enriched peptide samples were desalted as described before (Rappsilber et al., 2003).

Reversed-phase liquid chromatography-mass spectrometry (LC-MS) was performed using the UltiMate™ 3000 RSLCnano system (Dionex LC Packings/Thermo Fisher Scientific, Dreieich, Germany) coupled online to a Q Exactive Plus (Thermo Fisher Scientific, Bremen, Germany) instrument. The UHPLC system was equipped with two pre-columns (nanoEase™ M/Z Symmetry C18, 100 Å, 5 μm, Waters or μPAC™ trapping column, PharmaFluidics) and a corresponding analytical column (25 cm nanoEase™ M/Z HSS C18 T3 column, Waters or 50 cm μPAC™ column, PharmaFluidics). The MS instrument was externally calibrated using standard compounds and equipped with a nanoelectrospray ion source and a fused silica emitter (New Objectives). For MS analysis, dried peptides were resolved in 15 μl 0.1% TFA and analysed with an 1h LC method. Gradients were applied using a binary solvent systems of 0.1% FA (v/v, solvent A, ‘A’) and 0.1% FA/86% acetonitrile (v/v, solvent B, ‘B’). For nanoEase column setup, a gradient from 4% B to 42% B in 30 min and to 95% B in 3 min was performed, followed by a re-equilibration with 4% B for 16 min. μPAC columns were used with a gradient from 1-24% B in 22 min, followed by an increase to 42% B in 11 min and to 95% B in 6 min. Re-equilibration was performed with 1% B for 16 min. Full scans were acquired for a mass range of 375-1,700 *m/z* with a resolution of 70,000 at 200 *m/z*. The automatic gain control (AGC) was set to 3e^6^ ions with a max. ion time (IT) of 60 ms. MS/MS analyses of multiply charged peptide ions were generally performed using a top12 method and higher-energy collisional dissociation (HCD) with an energy of 28 and an exclusion time of 45 s. The resolution for MS/MS scans was 35,000 and the AGC 1e5 with a max. IT of 120 ms.

### Statistical analysis

Protein sequence alignment was performed by Clustal Omega (1.2.4) Multiple Sequence Alignment (Madeira et al., 2019). Immunoblot signals were quantified using ImageJ or GeneTools (Syngene) for film and CCD images, respectively. Two-tailed unpaired t-tests were used for statistical comparisons between 2 groups. For experiments containing more groups, one-way ANOVA with Dunnett’s post hoc test was used to determine statistical differences against a control mean, or one-way ANOVA with Tukey’s post hoc test was used to determine statistical difference between the mean of all possible pairs. Statistical analyses were performed on GraphPad Prism (v8.1.2). Data are presented as mean ± SEM. * p<0.05, ** p<0.01, *** p<0.001, **** p<0.0001.

### Bioinformatics

Peak lists obtained from MS/MS spectra were identified using Mascot version 2.6.1 [PMID 10612281] and MS Amanda version 2.0.0.9695 [PMID 24909410]. The search was conducted using SearchGUI version [3.3.17] [PMID 21337703]. Protein identification was conducted against a concatenated target/decoy [PMID 20013364] version of the Homo sapiens complement of the UniProtKB (version of 04/2019; 95,916 target sequences). The decoy sequences were created by reversing the target sequences in SearchGUI. The identification settings were as follows: trypsin, specific, with a maximum of 4 missed cleavages; thermolysin, unspecific; both with 5 ppm as MS1 and 0.5 Da as MS2 tolerances. Fixed modifications were set to: carbamidomethylation of C; variable modifications to: acetylation of protein N-term, phosphorylation of S and T, oxidation of M. All algorithm-specific settings are listed in the Certificate of Analysis available in the supplementary information. Peptides and proteins were inferred from the spectrum identification results using PeptideShaker version 1.16.44 [PMID 25574629]. Peptide Spectrum Matches (PSMs), peptides and proteins were validated at a 1% False Discovery Rate (FDR) estimated using the decoy hit distribution. Post-translational modification localizations were scored using the D-score [PMID 23307401] and the phosphoRS score [PMID 22073976] with a threshold of 95 as implemented in the compomics-utilities package [PMID 21385435]. A phosphoRS score > 95 was considered as a confident site localization.

## Data availability

All raw data and original Mascot result files have been deposited to the ProteomeXchange Consortium (http://proteomecentral.proteomexchange.org) via the PRIDE partner repository (http://www.ebi.ac.uk/pride/archive/login) (Perez-Riverol et al., 2019) with the dataset identifier PXD018005.

## Online Supplemental Material

Fig. S1 provides additional evidence to support the data shown in Fig. 1 that the ACBD5-VAPB binding is sensitive to phosphatase treatment. Fig. S2 gives an overview of the potential phosphorylation sites involved in the ACBD4/5-VAPB interaction explored in this study. Fig. S3 provides evidence that ACBD4 phosphomimetic mutants increase VAPB interaction and shows the subcellular localisation of ACBD4/5 mutants and their effect on peroxisomal morphology. Fig. S4 shows that expression of GSK3β results in increased cellular activity of the kinase, and that this affects peroxisome morphology in dMFF cells. Additionally, it explores the interaction of VAPB mMSP with ACBD5. Fig S5 shows that in addition to ACBD4 and ACBD5, other FFAT-containing proteins have a serine/threonine residue at position 5 of the FFAT core.

## Acknowledgements

We thank all colleagues who provided cell lines, plasmids, and antibodies (see Tables S1–S6 for more information), T.A. Schrader, A. Correia and P. Cherek for technical assistance and the PRIDE team for data deposition to the ProteomeXchange Consortium. This work was supported by grants from the Biotechnology and Biological Sciences Research Council (BB/N01541X/1 to M. Schrader; BB/T002255/1 to M. Schrader and J.L. Costello), Medical Research Council (CiC 08135 to M. Schrader), UKRI Future Leader Fellowship Award (MR/T019409/1 to J.Costello), Royal Society Research Grant Award (RGS\R2\192378 to J. Costello), University of Exeter, and the European Union’s Horizon 2020 research and innovation programme under the Marie Skłodowska-Curie grant agreement No 812968 PERICO (to M. Schrader and B. Warscheid) as well as the Deutsche Forschungsgemeinschaft (DFG, German Research Foundation) Project-ID 278002225/GRK 2202 (to B. Warscheid), FOR 1905 (PerTrans; to B. Warscheid), Project-ID 403222702 – SFB 1381 (to B. Warscheid) and under Germany’s Excellence Strategy – EXC-2189 – Project-ID 390939984 (CIBSS; to B. Warscheid). S. Kors is supported by the GW4 BioMed MRC Doctoral Training Partnership (MR/N0137941/1). Work included in this study has also been performed in partial fulfilment of the doctoral theses of R. Maier at the University of Freiburg. The research data supporting this publication are provided within this paper, as supplementary information, or are deposited on PRIDE.

## Author Contributions

C. Hacker, C. Bolton, R. Maier, L. Reimann, and E.J.A. Kitchener performed experiments and analyzed data. M. Schrader, J.L. Costello, B. Warscheid, and S. Kors conceived the project, performed experiments, and analyzed data. M. Schrader, J.L. Costello, and S. Kors wrote the manuscript, and all authors contributed to methods.

## Abbrevations

ACBD: acyl-coenzyme A-binding domain containing protein
ER: endoplasmic reticulum
FFAT: two phenylalanines (FF) in an acidic tract
λPP: λ protein phosphatase
GSK3β: glycogen synthase kinase-3 beta
IP: immunoprecipitation
MS: mass spectrometry
MSP: major sperm protein
VAP: vesicle-associated membrane protein (VAMP)-associated protein
VLCFA: very long-chain fatty acid.

## Supplementary Information

**Table S1.** Plasmids used in this study. Number indicates isoform.

| Plasmid | Source | Vector |
| --- | --- | --- |
| FLAG-ACBD4.2 | (Costello et al., 2017c) | pCMV-Tag2B |
| FLAG-ACBD5.1 | (Costello et al., 2017b) | pCMV-Tag2B |
| FLAG-ACBD5.1 mFFAT | (Costello et al., 2017b) | pCMV-Tag2B |
| Myc-ACBD5.1 | (Costello et al., 2017b) | pCMV-Tag3B |
| GSK3β | C. Miller, King's College London, England, UK | |
| Myc-VAPB | C. Miller, King's College London, England, UK | pCI-neo |
| GST-VAPB | (Costello et al., 2017b) | pGEX-6p2 |
| - | M. Wilmanns, EMBL Hamburg, Germany | pETM12 |

**Table S2.**
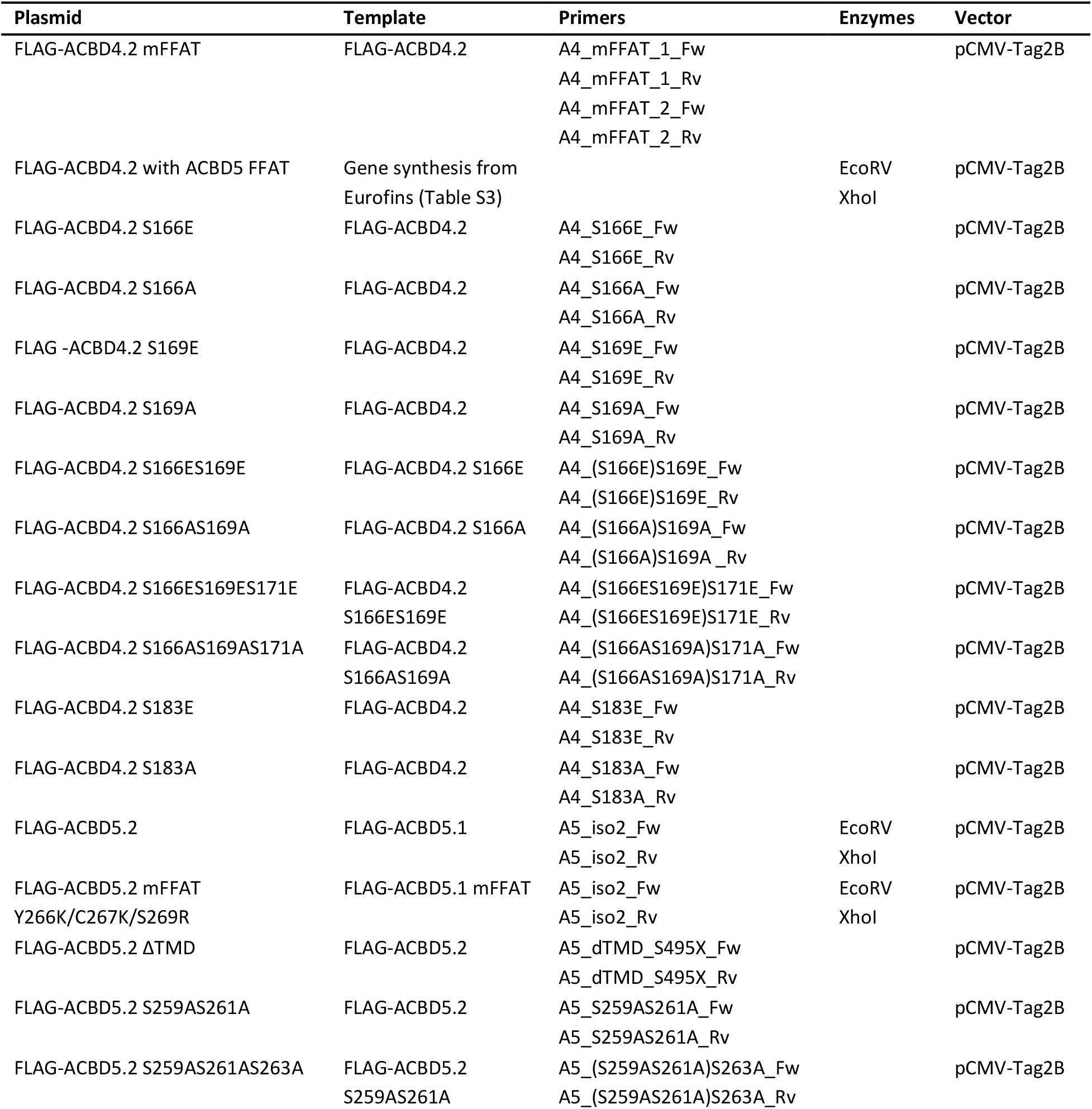
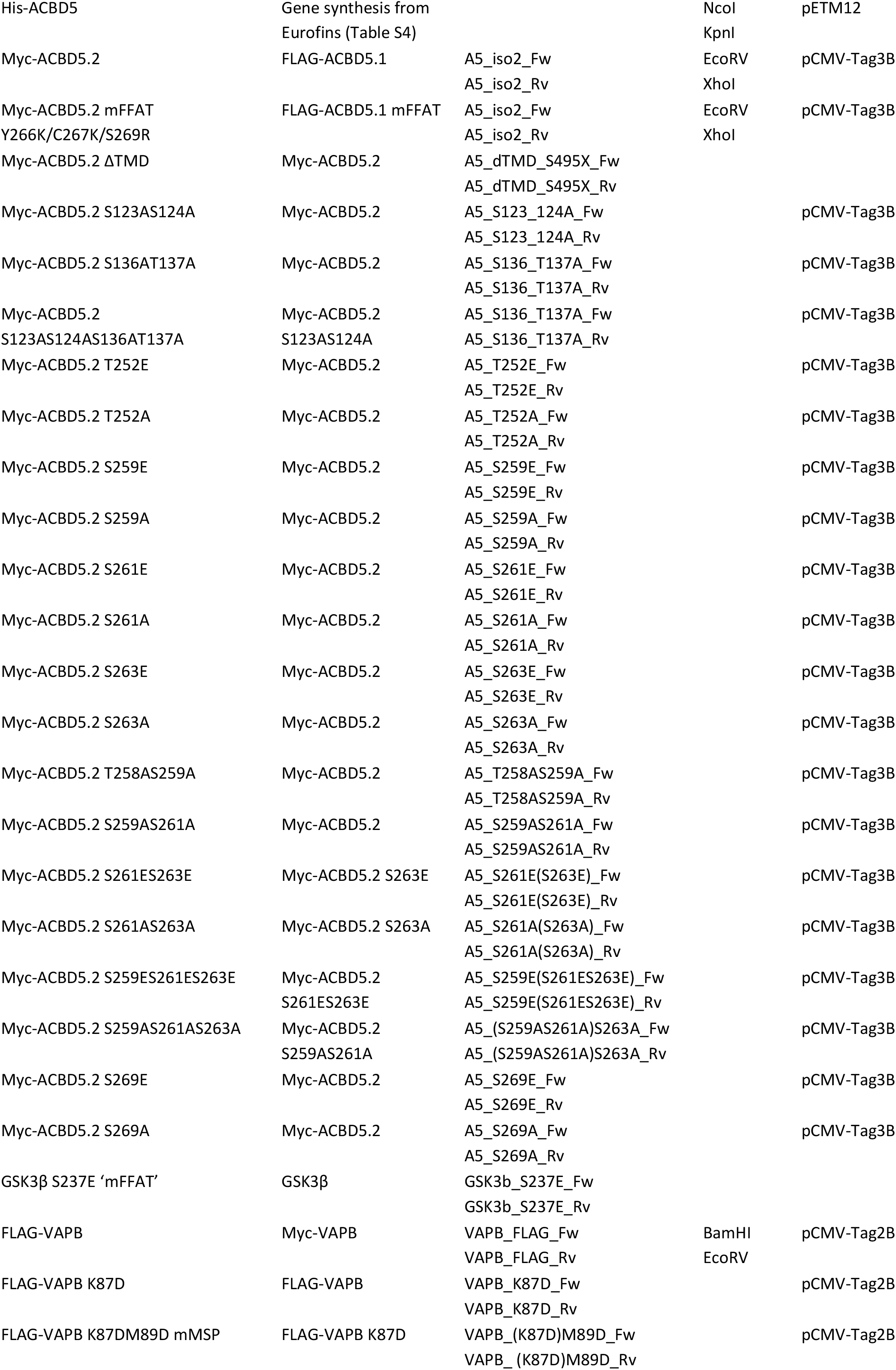
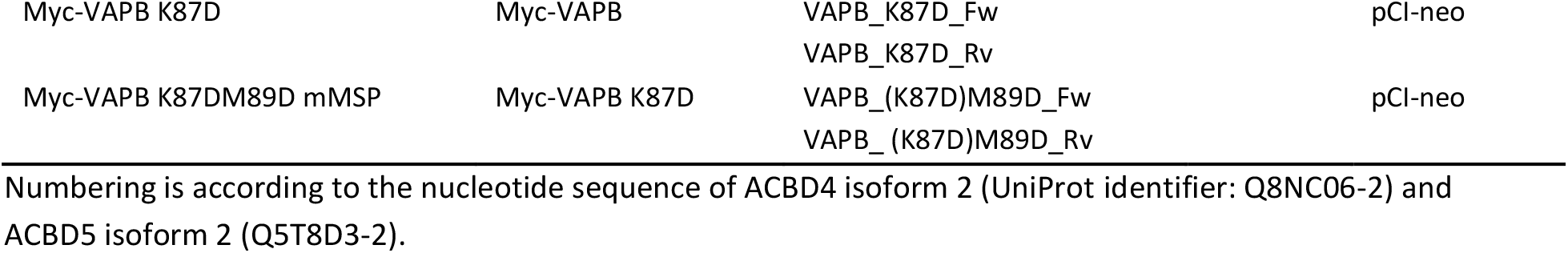
Plasmids generated in this study.

**Table S3.**
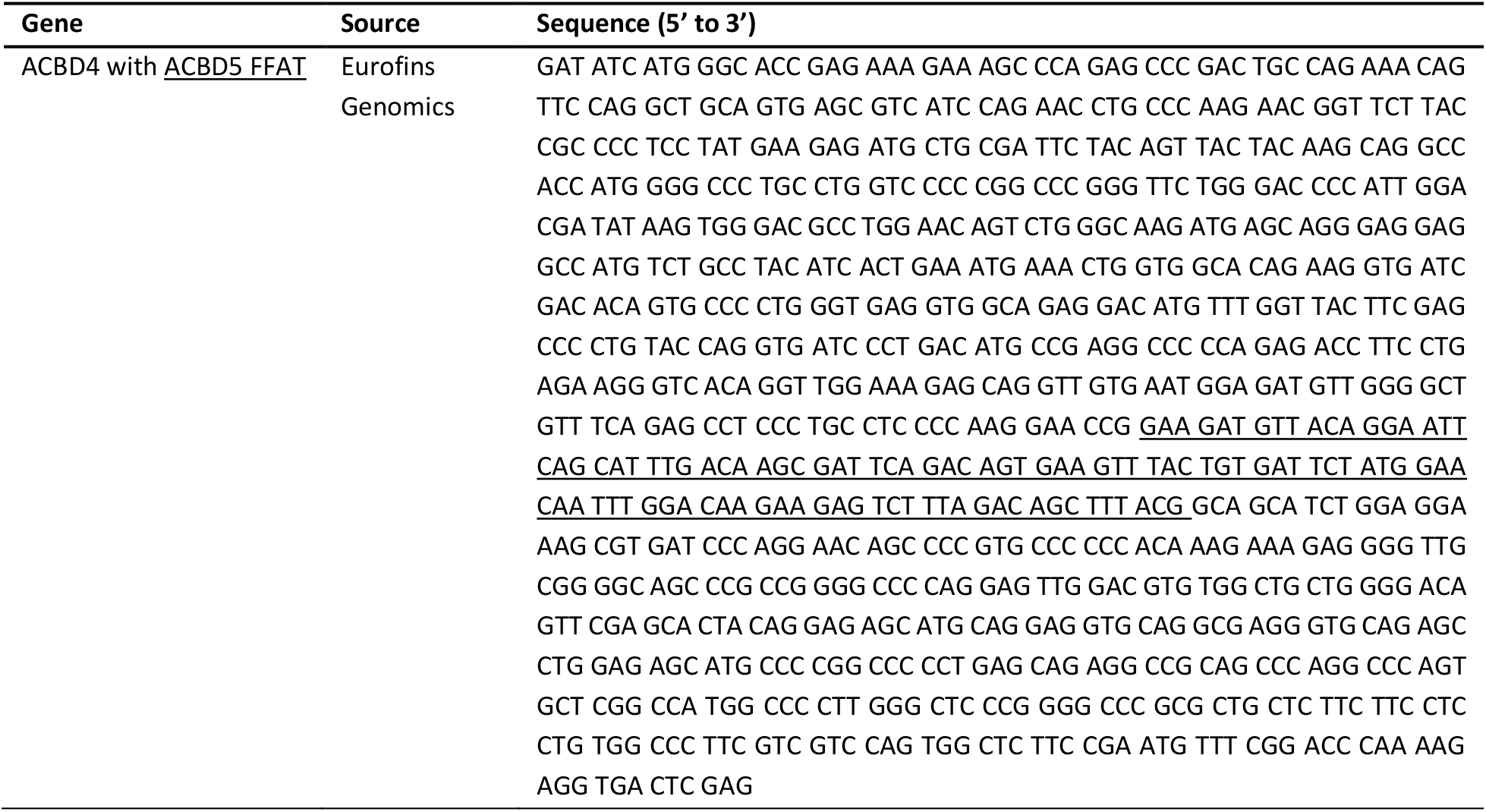
Sequence of ACBD4 with ACBD5 FFAT-like motif region.

**Table S4.**
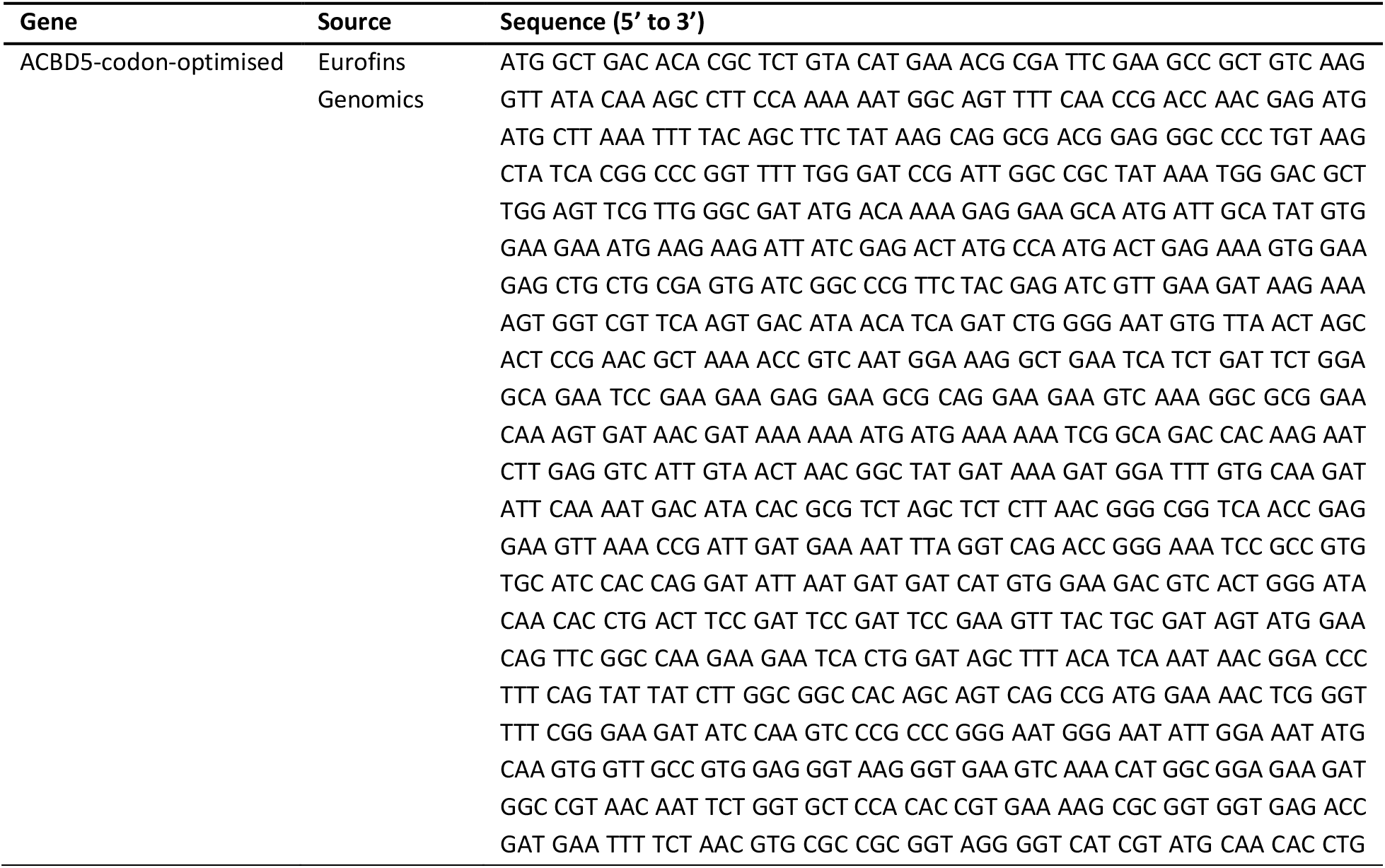
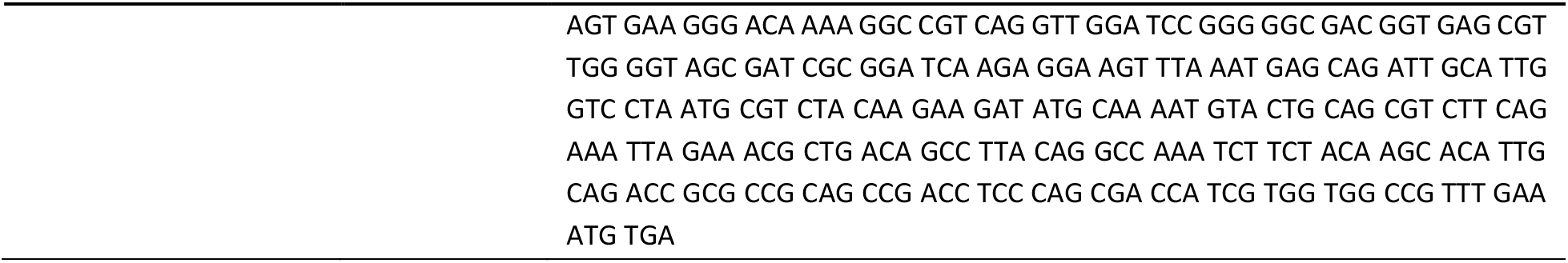
Condon optimised ACBD5 for expression in *E. coli*.

**Table S5.**
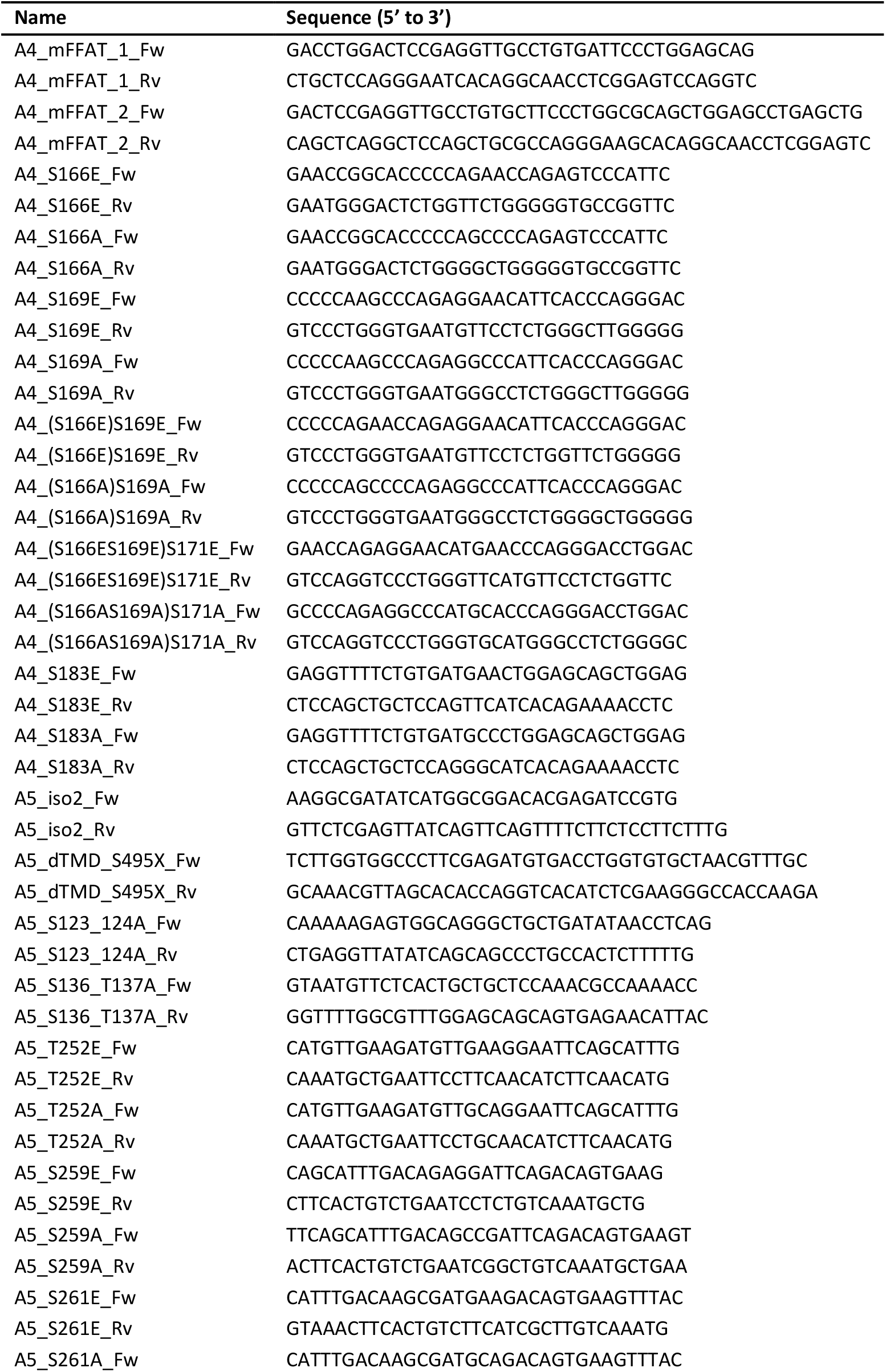
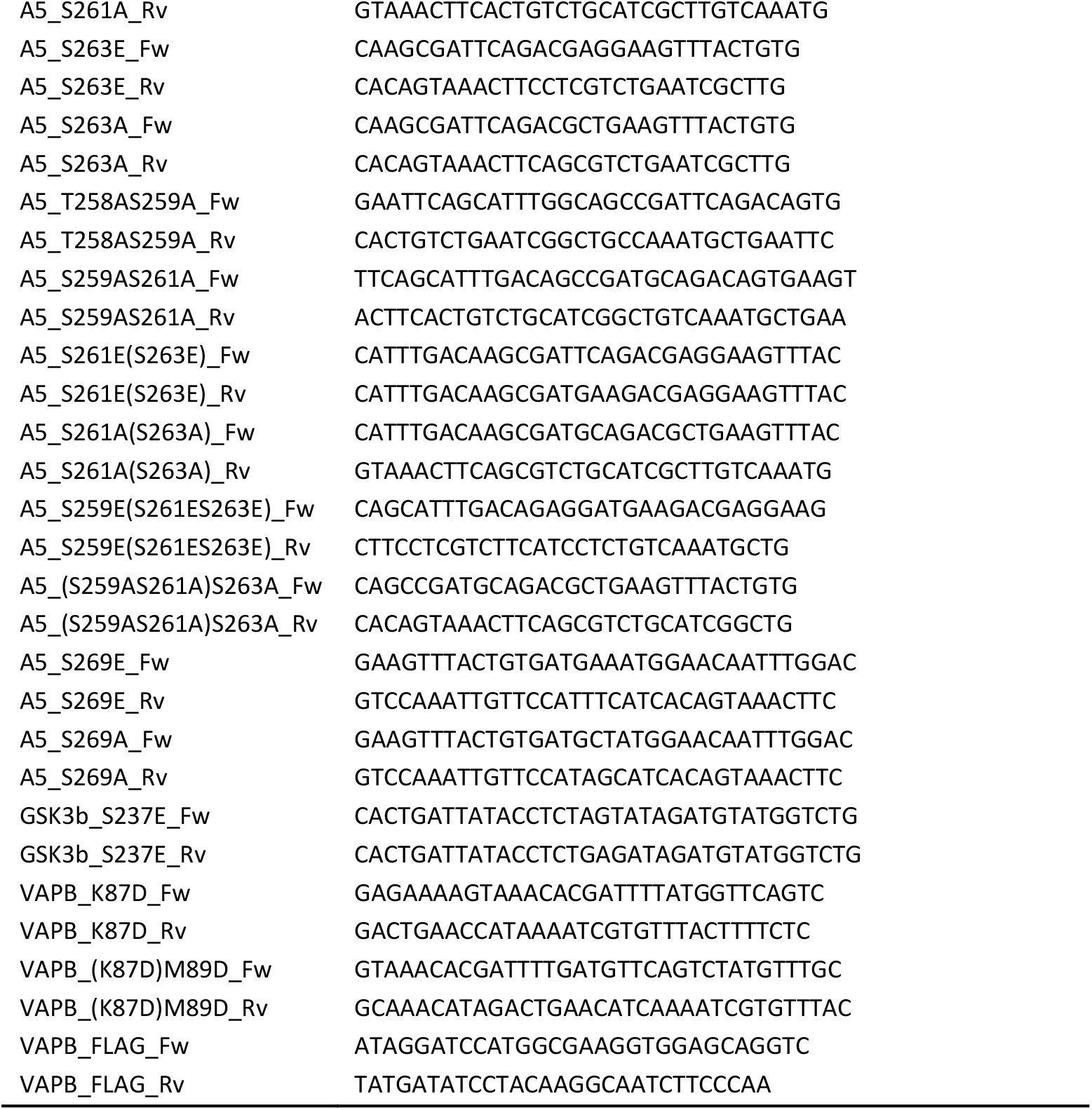
Primers used in this study.

**Table S6.**
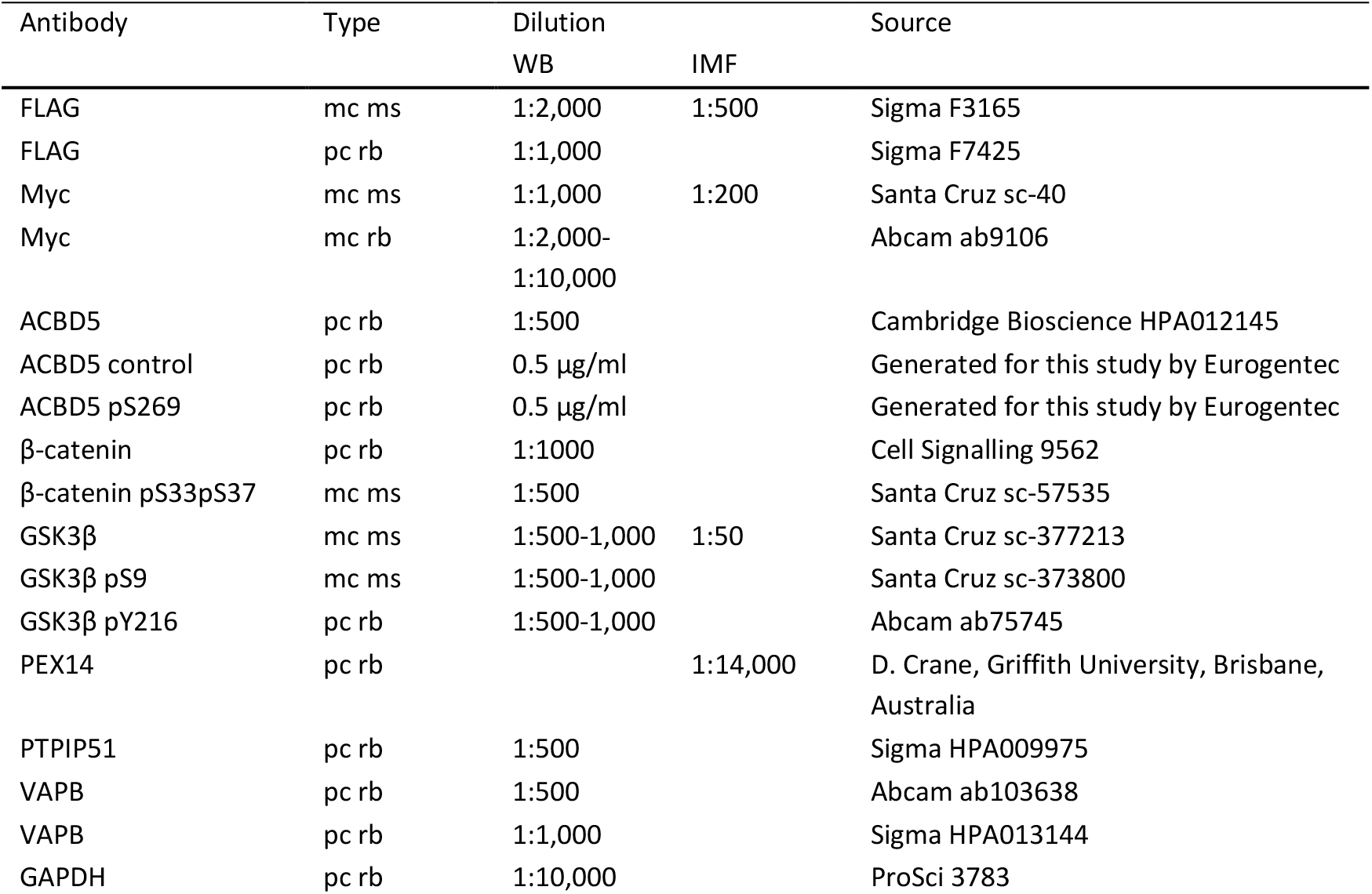
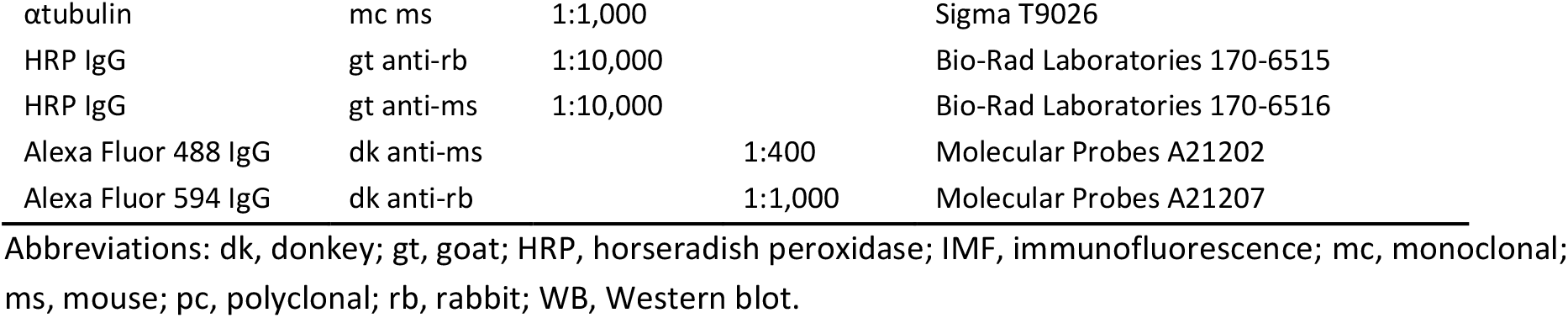
Primary and secondary antibodies used in this study.

## Figure Legends

**Supplementary Figure S1. The ACBD5-VAPB binding, but not the ACBD4-VAPB interaction, is sensitive to phosphatase treatment.** (**A**) FLAG-ACBD4/5 was expressed in COS-7 cells, and lysates were treated with or without λPP. FLAG-ACBD4/5 was immunoprecipitated and endogenous bound VAPB detected by immunoblotting using FLAG/VAPB antibodies. Constructs with mutations in the FFAT-like motif (mFFAT) were used as a negative control. (**B**) Myc-ACBD5 phospho-mutants were expressed in COS-7 cells and immunoprecipitated to detect endogenous bound VAPB using Myc/VAPB antibodies. WT, wild type.

**Supplementary Figure S2. Potential phosphorylation sites involved in the ACBD4/5-VAPB interaction.** (**A**) Schematic overview of replacement of the ACBD4 FFAT-like motif region (bold) by that of ACBD5. (**B**) Schematic overview of ACBD4 and ACBD5 domain structure, including the amino acid sequences of the FFAT-like motif region, with the phosphorylation sites mutated in this study in bold. (**C**) Alignment of the FFAT-like motif region of ACBD4 and ACBD5 from Hs, *Homo sapiens* (human); Rn, *Rattus norvegicus* (rat); Mm, *Mus musculus* (mouse); Cl, *Canis lupus familiaris* (dog); Fp, *Falco peregrinus* (Falcon); Xt, *Xenopus tropicalis* (frog); Dr, *Danio rerio* (zebrafish). Conservation of the phosphorylation sites reported in this study is indicated in bold. (**D**) Overview of the FFAT-like motif region of ACBD4 and ACBD5 and the constructs used in this study. Mutated residues are indicated in bold.

**Supplementary Figure S3. ACBD4 phosphomimetic mutants increase VAPB interaction.** (**A**) Subcellular localisation of ACBD4/5 constructs. COS-7 cells transfected with Myc-ACBD5 WT, mFFAT, S269E, S259A/S261A/S263A or ΔTMD; or FLAG-ACBD4 WT, S183E or S166A/S169A/S171A, were immunolabelled with PEX14 (peroxisomal marker) and Myc/FLAG antibodies. Bars: 10 μm (main), 2.5 μm (insets). (**B, C**) ACBD4 constructs with non-phosphorylatable (S → A) and phosphomimetic (S → E) residues upstream (S166/S169/S171) or within the FFAT core (S183) were generated and expressed in COS-7 cells. The FLAG-tagged proteins were immunoprecipitated and endogenous bound VAPB detected by immunoblotting using FLAG/VAPB antibodies. (**B**) Data were analyzed by one-way analysis of variance with Dunnett’s multiple comparison test. Total VAPB (IP fraction) was normalized against total VAPB (input) and FLAG-ACBD4 (IP fraction). Results of five independent IPs were quantified. (**D**) Quantification of peroxisome morphology in Myc-ACBD5 (S269E) or FLAG-ACBD4 (S183E) transfected COS-7 cells. Data were analyzed by one-way analysis of variance with Tukey’s multiple comparisons test. n = 400 per condition, from four replicates. ns, not significant; *, P < 0.05; ****, P < 0.0001. Error bars represent SEM. WT, wild type; mFFAT, mutated FFAT motif; TMD, transmembrane domain.

**Supplementary Figure S4. GSK3β expression alters peroxisome morphology in MFF-deficient fibroblasts.** (**A**) GSK3β expression increased phosphorylation of its substrate β-catenin, resulting in its degradation in COS-7 cells, as assessed by immunoblotting using GSK3β/GSK3β pY216/β-catenin/β-catenin pS33pS37 antibodies. GAPDH served as loading control. GSK3β K85A, catalytically inactive mutant; WT, wild type. (**B**) Myc-VAPB WT or mMSP (K87D/M89D), a mutant that cannot bind FFAT motifs, was co-expressed with FLAG-ACBD5 (or control vector (FLAG)). Myc-VAPB was immunoprecipitated and bound FLAG-ACBD5 detected by immunoblotting using Myc/FLAG antibodies. (C) Peroxisome morphology in MFF-deficient fibroblasts expressing GSK3β. Fixed cells were immunolabelled with PEX14 (peroxisomal marker) and GSK3β antibodies. Data were analyzed by a two-tailed unpaired t-test; ****, P < 0.0001. Error bars represent SEM. n = 800 per condition, from two independent experiments. Bars: 10 μm.

**Supplementary Figure S5. Examples of proteins with a serine/threonine residue at position 5 of the FFAT core.** A screen of human proteins with predicted FFAT motifs (score ≤2.5) identified by (Slee and Levine, 2019) revealed additional proteins, as well as ACBD4 and ACBD5, with a serine/threonine residue at position 5 of the FFAT core. These residues and (predicted) FFAT motifs showed conservation between species. Some of the serine/threonine residues at position 5 have been shown to be phosphorylated (indicated by an asterisk) (Hornbeck et al., 2015). The FFAT scores of the shown sequences are indicated (Murphy and Levine, 2016). FFAT motifs with a score of ≤3.5 are highlighted. Light green, acidic tract of the FFAT motif; dark green, FFAT core; orange, serine/threonine residue at position 5 of the FFAT core. Hs, *Homo sapiens* (human); Rn*, Rattus norvegicus* (rat); Mm, *Mus musculus* (mouse); Cl, *Canis lupus familiaris* (dog); Fp, *Falco peregrinus* (falcon); Xt, *Xenopus tropicalis* (frog); Dr, *Danio rerio* (zebrafish).

