## Supplemental Figures for "Regulating peroxisome-ER contacts via the ACBD5-VAPB tether by FFAT motif phosphorylation and GSK3β"

**Supplementary Figure S1**

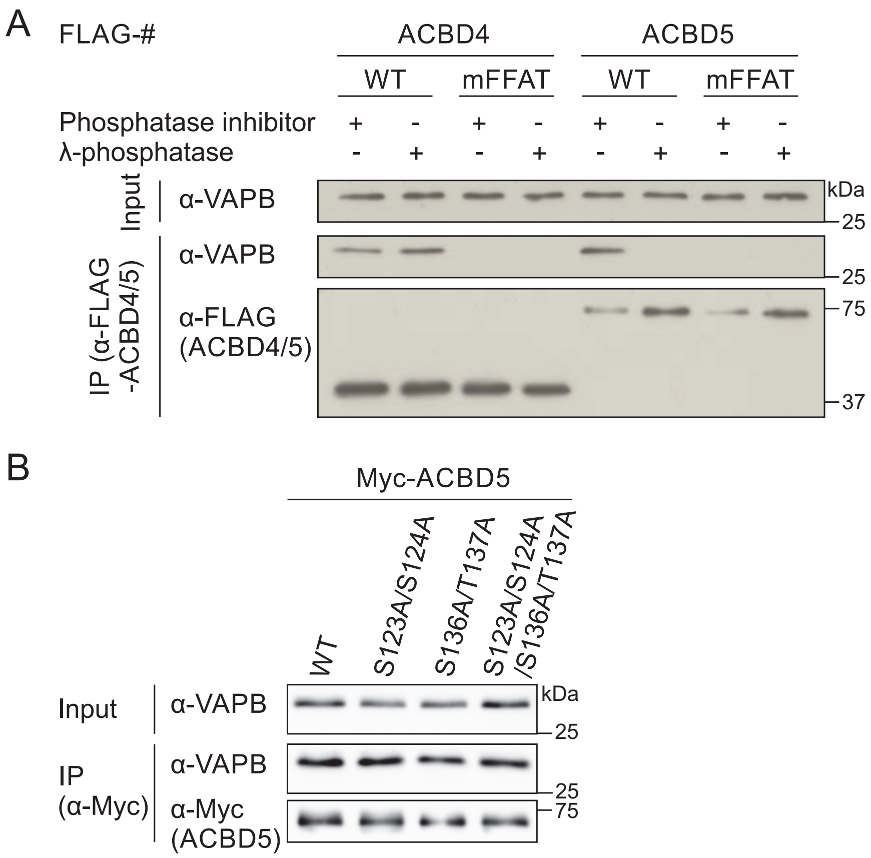

**Supplementary Figure S2**

A

| | | |
|---|---|---|
| ACBD5 | 243 | INDDHV**EDVTGIQHLTSDSDSEVYCDSMEQFGQEESLDSFT**SNNGPF 289 |
| ACBD4 | 157 | CLPKEP**APPSPESHSPRDLDSEVFCDSLEQLEPELVWTEQR**AASGGK 203 |
| ACBD4 with ACBD5 FFAT | 157 | CLPKEP**EDVTGIQHLTSDSDSEVYCDSMEQFGQEESLDSFT**AASGGK 203 |

B

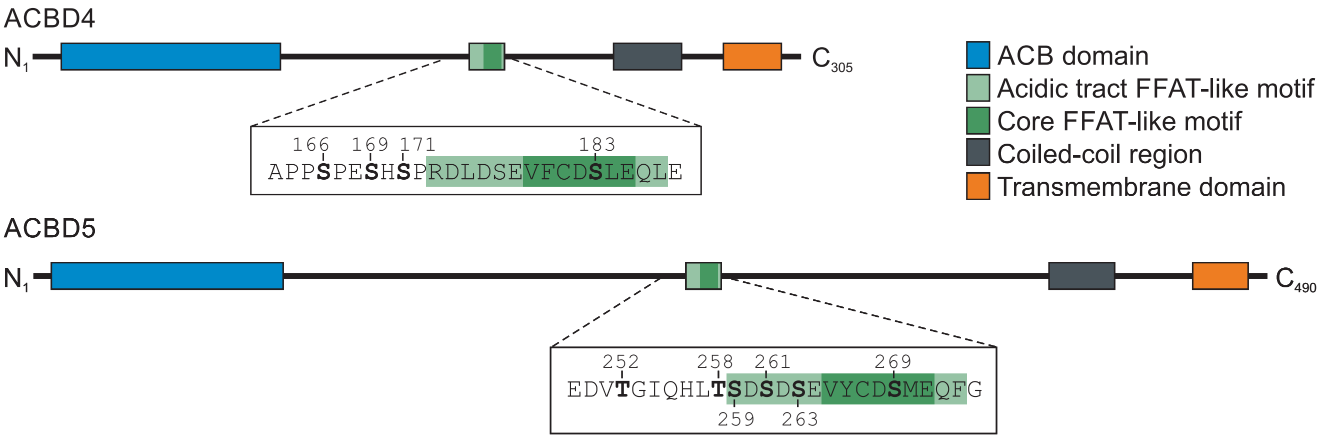


C

ACBD4 conservation (166 169 171 183)

| | |
|---|---|
| Hs | APP**S**PE**S**H**S**PRDLDSEVFCD**S**LEQLEPEL------ |
| Rn | APP**S**PE**S**RLPRDLDLEVFCD**S**VEQLEPELVRVPVL |
| Mm | VPP**S**PE**S**RPPRDLDLEVFCD**S**VEQLEPELVRLPVL |
| Cl | EPP**S**PE**S**QPPRDLDSEVFCD**S**LEQLEPEL------ |
| Fp | TEALEG**S**QVTSDSEGDVYCD**T**LEQMEPEQVSGSLV |
| Xt | TED**S**SD**S**LEQLDLDK-DFND**T**VTWIYPHLFRPSSD |
| Dr | QRM**S**ECQGPISDTESEVFCD**S**LEQMDNIKLAVPDN |

ACBD5 conservation (252 258/259/261/263 269)

| | |
|---|---|
| Hs | EDV**T**GIQHL**TS**D**S**D**S**EVYCD**S**MEQFGQEESLDSFT |
| Rn | EDA**S**GVHHL**TS**D**S**D**S**EVYCD**S**MEQFGQE-------- |
| Mm | EDA**S**GIHHL**TS**D**S**D**S**EVYCD**S**MEQFGQE-------- |
| Cl | EDV**S**GIQHL**TS**D**S**D**S**EVYCD**S**MEQFGQEESLDSFT |
| Fp | EEV**S**AAQHL**TS**D**S**D**S**EVYCD**S**MEQLGLEEPLEIIT |
| Xt | EDIAAMQHL**TS**D**S**D**S**EIFCD**S**MEQFGQDEAHSLLL |
| Dr | EDV**S**SSHHVA**S**D**S**D**S**EVYCD**S**VDQFGGEDGSEIHM |

D

ACBD4 constructs (166 169 171 183)

| | |
|---|---|
| WT | APP**S**PE**S**H**S**PRDLDSEVFCD**S**LEQLE |
| mFFAT | APPSPESHSPRDLDSEV**A**C**A**SL**A**QLE |
| S166E | APP**E**PESHSPRDLDSEVFCDSLEQLE |
| S166A | APP**A**PESHSPRDLDSEVFCDSLEQLE |
| S169E | APPSPE**E**HSPRDLDSEVFCDSLEQLE |
| S169A | APPSPE**A**HSPRDLDSEVFCDSLEQLE |
| S166E/S169E | APP**E**PE**E**HSPRDLDSEVFCDSLEQLE |
| S166A/S169A | APP**A**PE**A**HSPRDLDSEVFCDSLEQLE |
| S166E/S169E/S171E | APP**E**PE**E**H**E**PRDLDSEVFCDSLEQLE |
| S166A/S169A/S171A | APP**A**PE**A**H**A**PRDLDSEVFCDSLEQLE |
| S183E | APPSPESHSPRDLDSEVFCD**E**LEQLE |
| S183A | APPSPESHSPRDLDSEVFCD**A**LEQLE |

ACBD5 constructs (252 258/259/261/263 269)

| | |
|---|---|
| WT | EDV**T**GIQHL**TS**D**S**D**S**EVYCD**S**MEQFG |
| mFFAT | EDVTGIQHLTSDSDSEV**KK**D**K**MEQFG |
| T252E | EDV**E**GIQHLTSDSDSEVYCDSMEQFG |
| T252A | EDV**A**GIQHLTSDSDSEVYCDSMEQFG |
| S259E | EDVTGIQHLT**E**DSDSEVYCDSMEQFG |
| S259A | EDVTGIQHLT**A**DSDSEVYCDSMEQFG |
| S261E | EDVTGIQHLTSD**E**DSEVYCDSMEQFG |
| S261A | EDVTGIQHLTSD**A**DSEVYCDSMEQFG |
| S263E | EDVTGIQHLTSDSD**E**EVYCDSMEQFG |
| S263A | EDVTGIQHLTSDSD**A**EVYCDSMEQFG |
| T258E/S259E | EDVTGIQHL**EE**DSDSEVYCDSMEQFG |
| T258A/S259A | EDVTGIQHL**AA**DSDSEVYCDSMEQFG |
| S259E/S261E | EDVTGIQHLT**E**D**E**DSEVYCDSMEQFG |
| S259A/S261A | EDVTGIQHLT**A**D**A**DSEVYCDSMEQFG |
| S261E/S263E | EDVTGIQHLTSD**E**D**E**EVYCDSMEQFG |
| S261A/S263A | EDVTGIQHLTSD**A**D**A**EVYCDSMEQFG |
| S259E/S261E/S263E | EDVTGIQHLT**E**D**E**D**E**EVYCDSMEQFG |
| S259A/S261A/S263A | EDVTGIQHLT**A**D**A**D**A**EVYCDSMEQFG |
| S269E | EDVTGIQHLTSDSDSEVYCD**E**MEQFG |
| S269A | EDVTGIQHLTSDSDSEVYCD**A**MEQFG |

**Supplementary Figure S3**

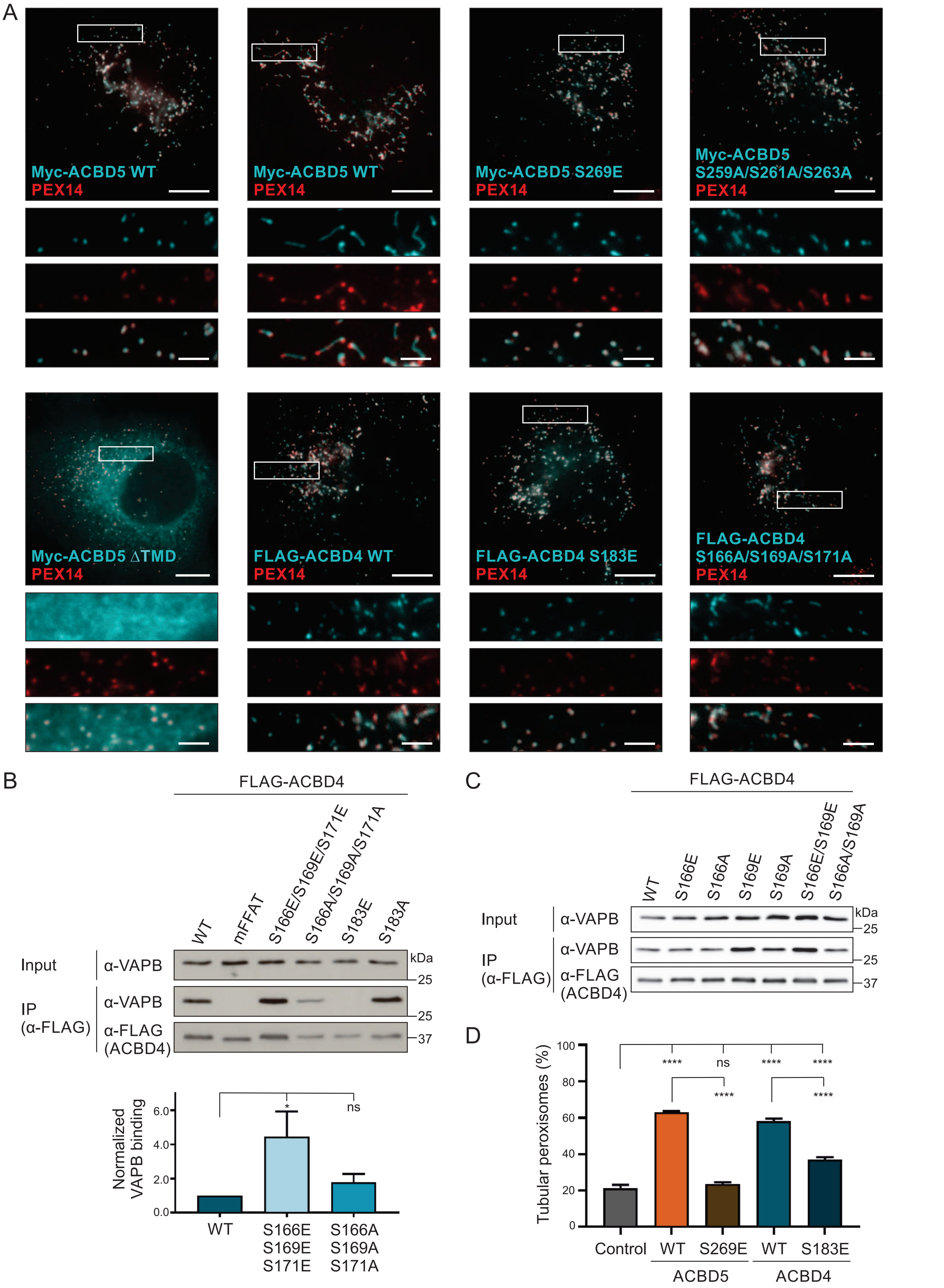

**Supplementary Figure S4**

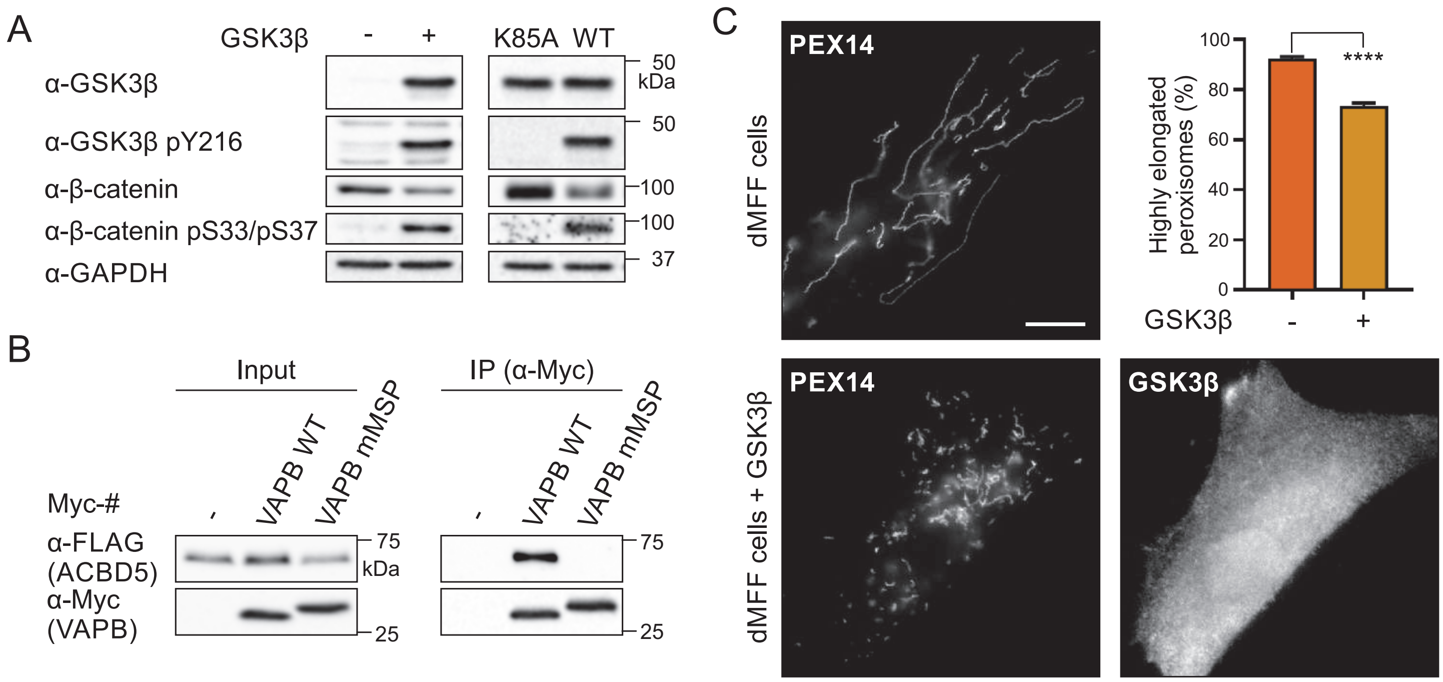

**Supplementary Figure S5. Examples of proteins with a serine/threonine residue at position 5 of the FFAT core.**

**ACBD4** – Acyl-CoA-binding domain-containing protein 4 (Q8NC06)

| | | |
|---|---|---|
| Hs | SHSPRDLDSEVFCDSLEQLEPELVWTE | **3.5** |
| Rn | SRLPRDLDSEVFCDSVEQLEPELVRVP | 3.5 |
| Mm | SRPPRDLDSEVFCDSVEQLEPELVRLP | 3.5 |
| Cl | SQPPRDLDSEVFCDSLEQLEPELVWTE | 3.5 |
| Fp | SQVTSDSEGDVYCDTLEQMEPEQVSGS | 2.5 |
| Xt | SLEQLDLDK-DFNDTVTWIYPHLFRPS | 3.5 |
| Dr | QGPISDTESEVFCDSLEQMDNIKLTAV | 2.5 |

**ACBD5** – Acyl-CoA-binding domain-containing protein 5 (Q5T8D3)

| | | | |
|---|---|---|---|
| Hs | QHLTSDSDSEVYCDSMEQFGQEESLDS | **2.5** | * |
| Rn | HHLTSDSDSEVYCDSMEQFGQEEYYLG | 2.5 | |
| Mm | HHLTSDSDSEVYCDSMEQFGQEEYYLG | 2.5 | |
| Cl | QHLTSDSDSEVYCDSMEQFGQEESLDS | 2.5 | |
| Fp | QHLTSDSDSEVYCDSMEQLGLEEPLEI | 2.5 | |
| Xt | QHLTSDSDSEIFCDSMEQFGQDEADHS | 2.5 | |
| Dr | HHVASDSDSEVYCDSVDQFGGEDGSEI | 2.5 | |

---

**CALCOCO1** – Calcium-binding and coiled-coil domain-containing protein 1 (Q9P1Z2)

CALCOCO1 has been shown to interact with VAP (Nthiga et al., 2020). Mutations in the FFAT motif strongly reduced the interaction.

| | | |
|---|---|---|
| Hs | DALEDHMDGHFFFSTQDPFTFE | 3.0 |
| Rn | DALEDHMDGHFFFSTQDPFTFE | 3.0 |
| Mm | DALEDHMDGHFFFSTQDPFTFE | 3.0 |
| Cl | DALEDHMDGHFFFSTQDPFTFE | 3.0 |
| Fp | QGFERHVQTHFDQNV---LNFD | 8.5 |
| Xt | ---------------------- | n/a |
| Dr | ---------------------- | n/a |

**SNX2** – Sorting nexin-2 (O60749)

SNX2 has been shown to interact with VAPB via two FFAT motifs (Dong et al., 2016). Mutating the phenylalanine (F2) in the FFAT motif below resulted in abolished VAPB binding. Mutating the residue in the other FFAT motif strongly reduced the interaction (DDREDLFAEATEEV – 2.0).

| | | |
|---|---|---|
| Hs | DFEDLEDGEDLFTSTVSTLESSPSSPE | **3.5** |
| Rn | DFEELEDGEDLFTSTVSTLESSPSSPE | 3.5 |
| Mm | DFEELEDGEDLFTSTVSTLESSPSSPE | 3.5 |
| Cl | DFAELEDGQDLFTSTVSTLESSPSSPE | 4.0 |
| Fp | --------------------------- | n/a |
| Xt | ECEDLEDGEDLFTSTVSTLESSPSSPE | 3.5 |
| Dr | EQEDSEAAEELFVSVM-------ESPE | 5.0 |

**AKAP11/AKAP220** – A-kinase anchor protein 11 (Q9UKA4)

A peptide containing the FFAT region of the human AKAP11 protein has been shown to bind Scs2p, the major VAP homolog in yeast (Mikitova and Levine, 2012).

```
Hs      DIEDSDSEVSEFFDSFDQFDELEQTLE    1.0
Rn      DIEDSDSEVSEFFDSFDQFDELEQTLE    1.0
Mm      DVEDSDSEVSEFFDSFDQFDELEQTLE    1.0
Cl      DIEDSDSEVSEFFDSFDQFDELEQTLE    1.0
Fp      GIEDSDSEVSEFFDSFDQFDELEQALE    1.0
Xt      DVEDSDSELSEFFDSFDQFDETEASLE    1.0
Dr      GVEDSDSEVSEFFDSFDQFDELDQSFD    1.0
```

**JMY** - Junction-mediating and -regulatory protein (Q8N9B5)

Whether the predicted FFAT motif is involved in the JMY-VAP interaction has not been confirmed yet, but JMY has been found to immunoprecipitate with VAPA (Schlüter et al., 2014).

```
Hs      VLFTETDDPEEYYESLSELRQKGYEEV    1.5     *
Rn      VLFTETDDPEEYYESLSELRQKGYEEV    1.5
Mm      VLFTETDDPEEYYESLSELRQKGYEEV    1.5
Cl      VLFTETDDPEEYYESLSELRQKGYEEV    1.5
Fp      VLFTETDDPEEYYESLSELRQKGYEEV    1.5
Xt      VLFADSDDPEEYYQSLSELRHKGYEEG    3.5
Dr      VLFPDSEDAEEYYESLSELRQKGYEDA    1.5
```

**MYOME** – Myomegalin (Q5VU43)

```
Hs      AGDDTEDTSTEFTDSIEEEAAHHSHQQ    1.5
Rn      AGDETEDTSTQFTDSIEEEAAHNSHQQ    2.5
Mm      AGDETEDTSTEFTDSIEEEAAHTSHQQ    1.5
Cl      AGEDTEDASTEFTDSIEEEAAHHNHGG    2.0
Fp      TGTDADDASSTFTYSIKEEAAHGVATQ    6.0
Xt      DNDVDEDSSSQFSDSIEDDTDYQSNGQ    2.5
Dr      EEEDEEDCNSEFAGSGEDEKRSKRTAQ    4.0
```

**ATG2B** - Autophagy-related protein 2 homolog B (Q96BY7)

```
Hs      EESGSEEETLQYFSTVDPNYRSRRKKK    2.0
Rn      EDSGSEEETLQYFSAVDPNYRSRRKKK    1.5
Mm      EDSGSEEETLQYFSAVDPNYRSRRKKK    1.5
Cl      EESGSEEETLQYFSTVDPNYRSRRKKK    2.0
Fp      DESGSEEETLQYYSTVDPNYRSRRRKK    2.5
Xt      DDSGSEEETMQHDYIMDSNYHCRRKKK    8.5
Dr      ---------------------------    n/a
```

**DYST** - Dystonin (Q03001)

```
Hs      NTGTDT-DSDDDFYDTPLFE-----------DDDHDSLL    2.0
Rn      DTATDS-DSDDYFYDTPLFE-----------DEDHDSLI    3.0
Mm      DTATDS-DSDDYFYDTPLFE-----------DEDHDSLI    3.0
Cl      VTETDT-DSEGDFYDTPLFE-----------DDDHDSLL    2.0
Fp      ---------------KPL---------------------    n/a
Xt      LTHTTSSNENYEFYSPEHVVKRACGTSQGHSDNDTSDLE    5.5
Dr      STDTEAVRGSQDLYLPSICDIQPEAISTKDKMSHDVSRT    7.0
```

**PCLO** - Protein piccolo (Q9Y6V0)

```
Hs        RHSWHDED-----DEAFDESPELKYRETKSQE      2.5
Rn        RHSWHDED-----DETFDESPELKFRETKSQE      2.0
Mm        RHSWHDED-----DETFDESPELKFRETKSQE      2.0      *
Cl        RHSWHDDD-----DDTFDDSPEPRYRETTSQD      2.0
Fp        RHSWHDDD-----DDNFDESPEPKYRETKSQD      2.5
Xt        RHSWHDDDEEDEDEETYDESPEPKHRETKSQE      2.0
Dr        YLSSHDQD-----DQVVQVIPVSEALEV---S      7.7
```

**CERT** - Ceramide transfer protein (Q9Y5P4)

It has been shown that CERT binds VAP via a FFAT motif (SLINEEEFFDAVEAA – 1.0) (Kawano et al., 2006). CERT has a potential second/non-canonical FFAT motif (Murphy and Levine, 2016). This motif has not been confirmed yet (but is predicted to be in an unstructured region – a characteristic of FFAT motifs).

```
Hs       RDKVVEDDED---------DFPTTRSDGDFLHST-NG    2.5
Rn       RDKVVEDDED---------DFPTTRSDGDFLHNT-NG    2.5
Mm       RDKVVEDDED---------DFPTTRSDGDFLHNT-NG    2.5
Cl       RDKVVEDDED---------DFPTTRSDGDFLHNT-NG    2.5
Fp       RDKGKIVTENSYLRNIQLFTHPTVRSDGDFVHNSNSS    8.0
Xt       RDKV-EDDED---------DFLHSHPNGDYIHSS-IG    4.5
Dr       RDKV-SDEEE---------DFPTLRPDADYLLNNNNS    4.5
```


Dong, R., Y. Saheki, S. Swarup, L. Lucast, J.W. Harper, and P. De Camilli. 2016. Endosome-ER Contacts Control Actin Nucleation and Retromer Function through VAP-Dependent Regulation of PI4P. *Cell*. 166:408–423. doi:10.1016/j.cell.2016.06.037.

Kawano, M., K. Kumagai, M. Nishijima, and K. Hanada. 2006. Efficient trafficking of ceramide from the endoplasmic reticulum to the golgi apparatus requires a VAMP-associated protein-interacting FFAT motif of CERT. *J. Biol. Chem.* 281:30279–30288. doi:10.1074/jbc.M605032200.

Mikitova, V., and T.P. Levine. 2012. Analysis of the key elements of FFAT-like motifs identifies new proteins that potentially bind VAP on the ER, including two AKAPs and FAPP2. *PLoS One*. 7:e30455. doi:10.1371/journal.pone.0030455.

Murphy, S.E., and T.P. Levine. 2016. VAP, a versatile access point for the endoplasmic reticulum: Review and analysis of FFAT-like motifs in the VAPome. *Biochim. Biophys. Acta - Mol. Cell Biol. Lipids*. 1861:952–961. doi:10.1016/j.bbalip.2016.02.009.

Nthiga, T.M., B. Kumar Shrestha, E. Sjøttem, J. Bruun, K. Bowitz Larsen, Z. Bhujabal, T. Lamark, and T. Johansen. 2020. CALCOCO 1 acts with VAMP - associated proteins to mediate ER -phagy. *EMBO J.* 39:e103649. doi:10.15252/embj.2019103649.

Schlüter, K., D. Waschbüsch, M. Anft, D. Hügging, S. Kind, J. Hänisch, G. Lakisic, A. Gautreau, A. Barnekow, and T.E.B. Stradal. 2014. JMY is involved in anterograde vesicle trafficking from the trans-Golgi network. *Eur. J. Cell Biol.* 93:194–204. doi:10.1016/j.ejcb.2014.06.001.
